# Regulatory Mechanisms Orchestrating Cellular Diversity in Cd36+ Olfactory Sensory Neurons Revealed by Single-Cell Multi-omics Analysis

**DOI:** 10.1101/2023.09.21.558403

**Authors:** Jiawen Yang, Peiyu Shi, Yiheng Li, Yachao Zuo, Tao Xu, Ziyang An, Dongjie Peng, Weixing Zhang, Yicong Xu, Zhongjie Tang, Anan Li, Jin Xu

## Abstract

The olfactory system relies on the precise expression of olfactory receptor (OR) genes in individual olfactory sensory neurons (OSNs) to detect and discriminate a vast array of odorants. Recent discoveries have revealed remarkable complexity and diversity within OSNs, including the existence of two distinct OSN populations based on high-affinity receptor Cd36 expression. However, the regulatory mechanisms governing this cellular diversity in the same cell type remain elusive.

To address these questions, we conducted single-cell multi-omics analyses of mature OSNs in the mouse olfactory epithelium. Firstly, we systematically revealed the transcriptome diversity and spatial distribution of Cd36+ OSNs and found a specific subset of olfactory receptors co-expressed with Cd36 in a deterministic manner. scATAC-seq profiling of chromatin landscape demonstrated a divergence between Cd36+ OSNs and Cd36- OSNs, including differential accessibility of cis-elements. By integrating transcriptome and epigenome profiling of OSN lineage-associated cell types, we revealed that the processes governing this diversity are initiated at the immature OSNs stage, where cellular diversity was first set by the lineage-specific binding of Lhx2 at Hdac9 enhancer. Hdac9, which is specifically expressed in the Cd36- OSN lineage, functions as a histone deacetylase and may repress the transcription of Mef2-dependent genes that contribute to Cd36+ OSN diversity. By gene regulation network analysis, we revealed Mef2a and Tshz1 as the key transcription factors, orchestrating the transcriptome diversity of Cd36+ OSNs. Remarkably, we identified and confirmed Tshz1 as a critical transcription factor that directly promotes Cd36 expression in OSNs through enhancer binding. Our study unravels the intricate regulatory landscape and principles governing cellular diversity in the olfactory system. These findings provide valuable insights into the regulation principles underlying neuronal heterogeneity and its functional implications.

## Introduction

The olfactory system plays a critical role in detecting and discriminating a wide range of odorants. Within the mouse olfactory epithelium (OE), individual olfactory sensory neurons selectively express one out of over 1,000 olfactory receptor (OR) genes (Buck and Axel, 1991; Chess et al., 1994), resulting in the remarkable “one-neuron-one-receptor” organization (Serizawa et al., 2003). The specific OR gene expression process not only determines odor sensitivity, but also bestows each neuron with a unique identity and activity pattern (Horgue et al., 2022; Nakashima et al., 2013; Serizawa et al., 2006; Tsukahara et al., 2021; Williams et al., 2011), guiding their axons to converge onto specific glomeruli within the olfactory bulb (OB) (Barnea et al., 2004; Imai et al., 2006; Mombaerts et al., 1996; Wang et al., 1998). Recent breakthroughs in systematic single-cell transcriptome analysis have unveiled a new dimension of complexity and diversity within the olfactory system (Brann et al., 2020; Finlay et al., 2022; Horgue et al., 2022; Shayya et al., 2022; Tsukahara et al., 2021; Vihani et al., 2020; Wang et al., 2022; Wu et al., 2018), Cd36, a high-affinity receptor with binary expression marked the existence of two distinct OSN populations in mouse (Horgue et al., 2022; Tsukahara et al., 2021; Wang et al., 2022). As Cd36 binds to various ligands, including the matrix protein thrombospondin (Dawson et al., 1997), long-chain fatty acids (FA) (Koonen et al., 2005), oxidized phospholipids (Podrez et al., 2002) and lipoproteins (Xu et al., 2021), it has been thought to play crucial roles in lipid odor recognition (Febbraio et al., 2001; Silverstein and Febbraio, 2009). Studies on CD36-deficient mice have revealed impairments in lipid mixture odors preference, and a drastic reduction in the number of OSNs responding to oleic acid, a major milk component (Oberland et al., 2015; Xavier et al., 2016).

Despite the preliminary descriptions of transcriptome diversity and the function of the Cd36-expressing OSNs, the regulatory mechanisms underlying this distinguished transcriptome diversity remain unclear. Several key questions pertaining to the binary expression of Cd36 within the OSN population remain unrevealed: What determines whether Cd36 is expressed or not within specific OSNs? Is there a coordination or interdependence between Cd36 expression and the selection of OR genes in individual OSNs? How and when does the expression of Cd36 becomes divergent and lead to the existence of two distinct OSN populations? These critical gaps in our understanding of the intricate molecular mechanisms governing cellular diversity within the olfactory system call for rigorous investigation.

In this study, we employed single-cell multi-omics approaches to comprehensively characterize the transcriptome and epigenome profiles of mature OSNs in the mouse main olfactory epithelium (MOE). By integrating transcriptome and epigenome data, we transcended the limitations of single-cell RNA sequencing alone and obtained a comprehensive view of the regulatory landscape underlying Cd36+ OSNs diversity. Through systematic integration analysis, we unveiled a deterministic regulatory mechanism of Cd36+ OSNs diversity, shedding light on the complex interplay between genetic and epigenetic factors.

The olfactory sensory neurons present an excellent model for investigating the basic regulatory principles of cellular diversity. Elucidating how the cellular diversity of Cd36+ OSNs is encoded and programmed using a multi-omics approach, will not only deepen our comprehension of the intricate cellular diversity within the olfactory system but also provide valuable insights into the broader understanding of neuronal heterogeneity and its functional implications.

## Results

### Transcriptome diversity and spatial distribution of Cd36+ olfactory sensory neurons

A previous study has demonstrated that the cellular diversity of the mouse OSN population can be decomposed into both cell identity-associated and activity-related gene expression modules, cellular identity denotes their spatial position in the main olfactory epithelium (MOE) and the expression of the lipid receptor (Tsukahara et al., 2021). Cd36+ OSNs, exhibit a significantly distinct transcriptome profile compared to Cd36- OSNs (Horgue et al., 2022; Tsukahara et al., 2021; Wang et al., 2022). We first reproduced and confirmed the transcriptome profile of the mouse MOE by performing single-cell RNA sequencing on 36,693 cells, with or without FACS enrichment for mature OSNs (**Methods; Figures S1A-S1E**). Unsupervised clustering of the mature OSN population revealed a similar distance and proportion of Cd36+ OSNs as reported in the study by Tsukahara et al (Tsukahara et al., 2021) (**Figure S2A; Figure 1A**). To increase the statistical power, we integrated the OSNs from these two datasets (**Methods; Figures S2B and S2C**). Among a total of 48,571 OSNs, 5.5% were Cd36+ OSNs (color in green), while 94.5% were Cd36- OSNs (color in gray) (**Figure 1A**).

**Figure 1.**
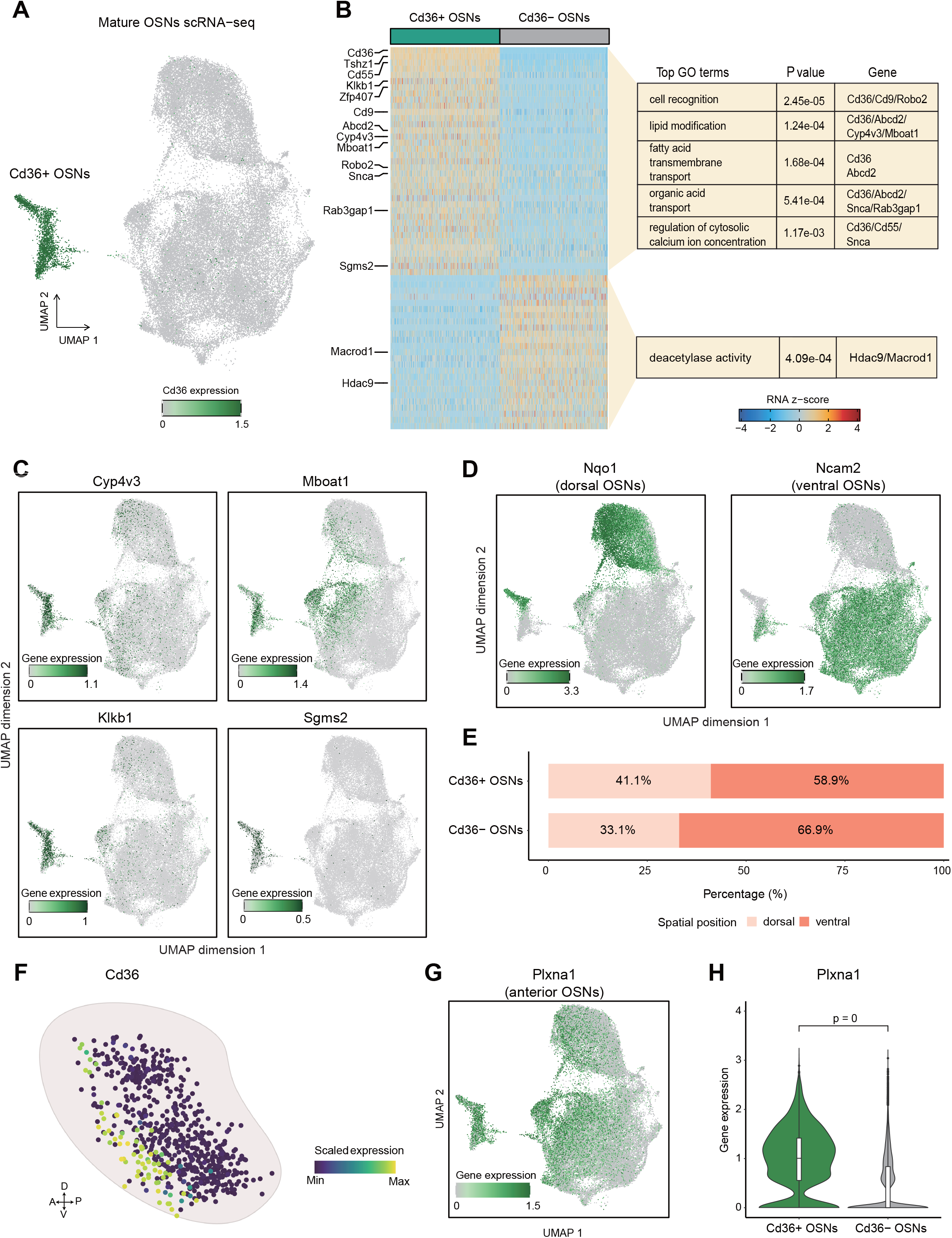
Transcriptome diversity of Cd36+ OSNs. (A) UMAP plot showing the subclustering result of the scRNA-seq data from mOSNs. Each cell is colored by its expression of Cd36. The cluster with high expression of Cd36 is annotated as Cd36+ OSNs. (B) Left: heatmap showing expression of differentially expressed genes (Cd36+ OSNs vs Cd36- OSNs, Wilcoxon Rank Sum test, Bonferroni adjusted p-value < 0.05, the absolute value of fold change >=1.5) in 500 cells sampling from Cd36+ OSNs and Cd36- OSNs, respectively. Each row represents a DEG, and each column represents a sampled cell. The color indicates the z-transformed gene expression (z-score range is - 4 to 4). Right: Top GO terms enriched for upregulated genes in Cd36+ OSNs and Cd36- OSNs (one-sided Fisher’s exact test with Benjamini-Hochberg correction). (C) UMAP plots visualizing the expression of upregulated genes in Cd36+ OSNs (Cyp4v3, Mboat1, Klkb1, and Sgms2). The minimum and maximum gene expression levels are shown on the bottom left of each panel. (D) UMAP plots visualizing the expression of markers for dorsal and ventral OSNs (Nqo1: a marker for dorsal OSNs, Ncam2: a marker for ventral OSNs). (E) Bar plot showing the percentage of dorsal and ventral cells in Cd36+ OSNs and Cd36- OSNs, respectively. (F) Scaled expression of Cd36 in olfactory glomeruli obtained from Wang et al(Wang et al., 2022). Schematic representation of the positions of the glomeruli in the olfactory bulb, which are identified or predicted by the spatial transcriptomics approach, scRNA-seq data, and a ridge regression model in Wang et al (Wang et al., 2022). Each dot represents an OR-associated glomerulus. (G) UMAP plot showing the expression of Plxna1 from scRNA-seq data. (H) Violin plot showing the expression of Plxna1 from scRNA-seq data. The expression levels of Plxna1 between Cd36+ OSNs and Cd36- OSNs are compared by the Wilcoxon Rank Sum test and the Bonferroni adjusted p-value is annotated above.

Next, we performed a comparative analysis of the transcriptome between Cd36+ and Cd36- OSNs (**Methods and Table S1**). We found that the Cd36+ OSNs-specific genes are functionally enriched in lipid modification, fatty acid transmembrane transport, and organic acid transport (**Figures 1B and 1C**), with representative genes shown in Figure 1C. The transcriptome diversity observed in Cd36+ OSNs further supports the previous functional study using Cd36-deficient mice, which has revealed that Cd36 may facilitate the detection of lipid odors (Oberland et al., 2015; Xavier et al., 2016). In addition to these gene ontology (GO) terms, we also identified an enrichment of upregulated genes in Cd36+ OSNs involved in the regulation of cytosolic calcium ion concentration. Previous studies have shown that CD36 on taste bud cells interacts with the fatty acids (FA) to induce intracellular calcium release from the ER (El-Yassimi et al., 2008). This calcium release then triggers calcium flux from membrane store-operated calcium (SOC) channels, ultimately leading to the release of neurotransmitter (Pepino et al., 2014). It suggests that Cd36, as a lipid-binding receptor, appears to play important roles in signaling perception and transduction in olfaction, as well as taste. Alongside genes associated with functional pathways, we identified two transcription factors, teashirt zinc finger family member 1 (Tshz1) and Zfp407, which are specifically expressed in Cd36+ OSNs (**Figure 1B; Figure S2D**). Conversely, we observed fewer downregulated genes in Cd36+ OSNs, showing no functional enrichment except for deacetylase activity (**Figure 1B**). One representative gene is Hdac9, which encodes histone deacetylase.

The spatial organization of OSNs in the OE can be divided into different zones or regions (Miyamichi et al., 2005; Tan and Xie, 2018; Zapiec and Mombaerts, 2020) and linked to the dorsoventral projection of their axons into specific regions of the olfactory bulb (Bashkirova et al., 2023). We then investigated how the Cd36 expression coordinates with the OSN spatial identity. Interestingly, we discovered a similar spatial distribution in Cd36+ and Cd36- OSNs along the dorsoventral axis, with a slightly higher ratio of dorsal to ventral in Cd36+ OSNs compared to Cd36- OSNs (**Figures 1D and 1E**). As odorant recognition is strongly tied to glomerular activity in the olfactory bulb (OB), we next examined the projections of Cd36+ OSNs onto the OB using the olfactory glomerular map obtained from Wang et al (Wang et al., 2022). This map, comprising 654 glomeruli, was constructed by profiling the positions of 65 glomeruli and training a linear model to predict the locations of other glomeruli based on transcriptional profiles of specific OSN types. Our analysis revealed that Cd36+ OSNs solely project to the anterior regions of the olfactory bulb based on the Cd36 expression (**Figure 1F**). This observation aligns with the previous immunostaining results, which showed the absence of CD36+ glomeruli in the posterior regions of the OB (Oberland et al., 2015). Furthermore, our scRNA-seq data revealed significantly higher expression of Plxna1, a well-studied marker gene for anterior projection (Imai et al., 2006; Nakashima et al., 2013; Sakano, 2010), in Cd36+ OSNs compared to Cd36- OSNs (**Figures 1G and 1H**). In summary, the unique molecular characteristics of Cd36+ OSNs mirror their function in lipid odor detection, which is independently regulated with dorsoventral position, while associated with their anterior projection.

### Cd36 co-expression with a subset of olfactory receptors

In the olfactory system, each OSN expresses a single OR allele from a large family of over 1000 genes, enabling the detection and discrimination of various odorants. We next wonder whether there exists a coordination or interdependence between Cd36 expression and the selection of OR genes in individual OSNs. By examining the cell counts of Cd36+ and Cd36- OSNs in each OSN subgroup, we evaluated whether there is a set of ORs specifically expressed or skewed towards one of the two groups (**Methods**). Our analysis revealed that out of 945 OSN groups containing at least 10 cells, 79 ORs exhibited significantly preferential expression in Cd36+ OSNs, while 90 ORs showed depleted expression in Cd36+ OSNs (**Figure S3A; Figure 2A**). For instance, Olfr806 and Olfr553 were identified as Cd36+ OSN-enriched OR genes, whereas Olfr536 and Olfr728 were found to be depleted in Cd36+ OSNs (**Figure 2B**). These results indicate that the expression of Cd36 may be associated with the OR choice.

To gain a better understanding of the ORs specifically expressed in Cd36+ OSNs, we examined their genomic locations and found a strong clustering of these ORs at the short arm of chromosome 10 (**Figures 2C and 2D**). Furthermore, the phylogenetic analysis demonstrated that these Cd36+ OSN-enriched ORs are distributed across a few clades, exhibiting lower sequence divergence within each clade (**Figure S3B**). OSNs with Cd36+ OSN-enriched ORs also project their axons into the anterior region of the OB (**Figure 2E**). In summary, these results indicate that a specific subset of ORs with similar characteristics co-expressed with Cd36 in a deterministic manner rather than a stochastic manner.

**Figure 2.**
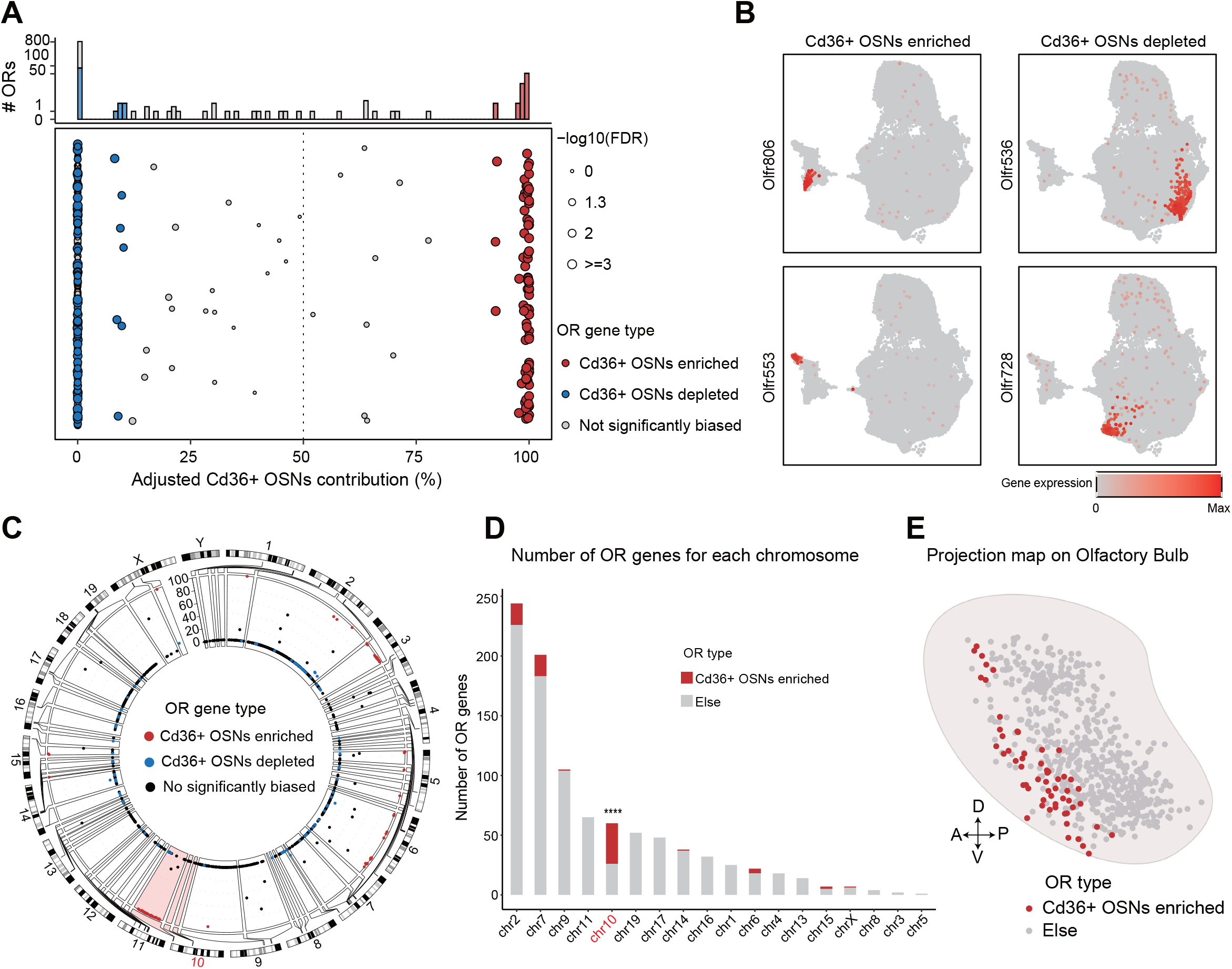
Cd36+ OSNs enriched ORs cluster in specific regions of the genome. (A) Bottom: dot plot visualizing the percentage of Cd36+ OSNs for each OSN subgroup that expresses a single OR gene. The percentage is adjusted by the number of cells in Cd36+ OSNs and Cd36- OSNs. Each dot represents an OR-associated OSN group, and it is assigned to one of three categories (Cd36+ OSNs enriched, Cd36+ OSNs depleted, and not significantly biased) using a binomial test. The dot color represents three categories of OR gene type and the dot size represents -log10 of Benjamini-Hochberg corrected p-value (FDR). Top: stacked bar plot showing the number of OSN subgroups for different levels of adjusted Cd36+ OSNs contributions. The color of the bar chart indicates the categories of OR gene type. (B) UMAP plot showing the expression of typical OR genes that are assigned to Cd36+ OSNs enriched OR genes and Cd36+ OSNs depleted OR genes, respectively. (C) Circular plot showing the genomic position for each OR gene. Tracks from outside to inside are the genome positions by chromosomes and the adjusted percentage of cells from Cd36+ OSNs. Each dot on the inner track represents an OR gene, and its color represents an OR gene type. The inner track within chromosome 10 is colored light red. (D) Stacked bar plot showing the number of OR genes within each chromosome. The color of the bar chart indicates if an OR gene is biased to Cd36+ OSNs. Chromosome enrichment analysis of Cd36+ OSN-enriched OR genes was performed by a binomial test. (E) Schematic representation of the positions of the glomeruli in the olfactory bulb. Each dot represents an OR-associated glomerulus, and its color indicates whether its associated OR gene is enriched in Cd36+ OSNs.

### Identification of transcription factors associated with chromatin divergence in Cd36+ OSNs

With a deeper understanding of the molecular diversity exhibited by Cd36+ OSNs, our next inquiry focused on the mechanism underlying this diversity. Single-cell chromatin accessibility profiling enables us to investigate the epigenomic landscape of OSNs at single-cell resolution. We obtained a chromatin accessibility profile of 13,158 cells from the mice olfactory epithelium, with FACS enrichment for mature OSNs (**Methods and Figures S4A-S4G**). Unsupervised clustering of scATAC-seq data for mature OSNs demonstrated a comparable divergence between Cd36+ and Cd36- OSNs at the epigenomic level, confirming that transcriptional diversity is encoded within the chromatin landscape (**Figure 3A; Figures S5A and S5B**). We explored the differences in accessible regions (or peaks) between Cd36+ and Cd36- OSNs and identified 311 up-regulated peaks and 181 down-regulated peaks in Cd36+ OSNs (**Figure 3B; Table S2**). Motif enrichment analysis of these differentially accessible peaks revealed potential transcription factors that may drive the chromatin diversity (**Figure 3B; Figures S5C and S5D**). Notably, we identified the binding motif of the myocyte enhancer factor 2 (MEF2) family, Foxd3, Stat2 and Foxj3 were enriched in the up-regulated peaks, while binding motif of Pknox2, Pou6f2, Meis2, Pbx3 and Lhx2 were enriched in the down-regulated peaks of Cd36+ OSNs (**Figure 3B; Figures S5C and S5D**). To further depict which gene in a protein family is the functional one in the regulation of Cd36+ OSN divergence, we compared the expression differences of transcription factors versus their binding motif accessibility. Intriguingly, except for Tshz1, most of the transcription factors that showed motif enrichment in the differential peaks demonstrated strong divergences in chromatin accessibility, but displayed barely any differences in expression levels (**Figures 3C and 3D**).

**Figure 3.**
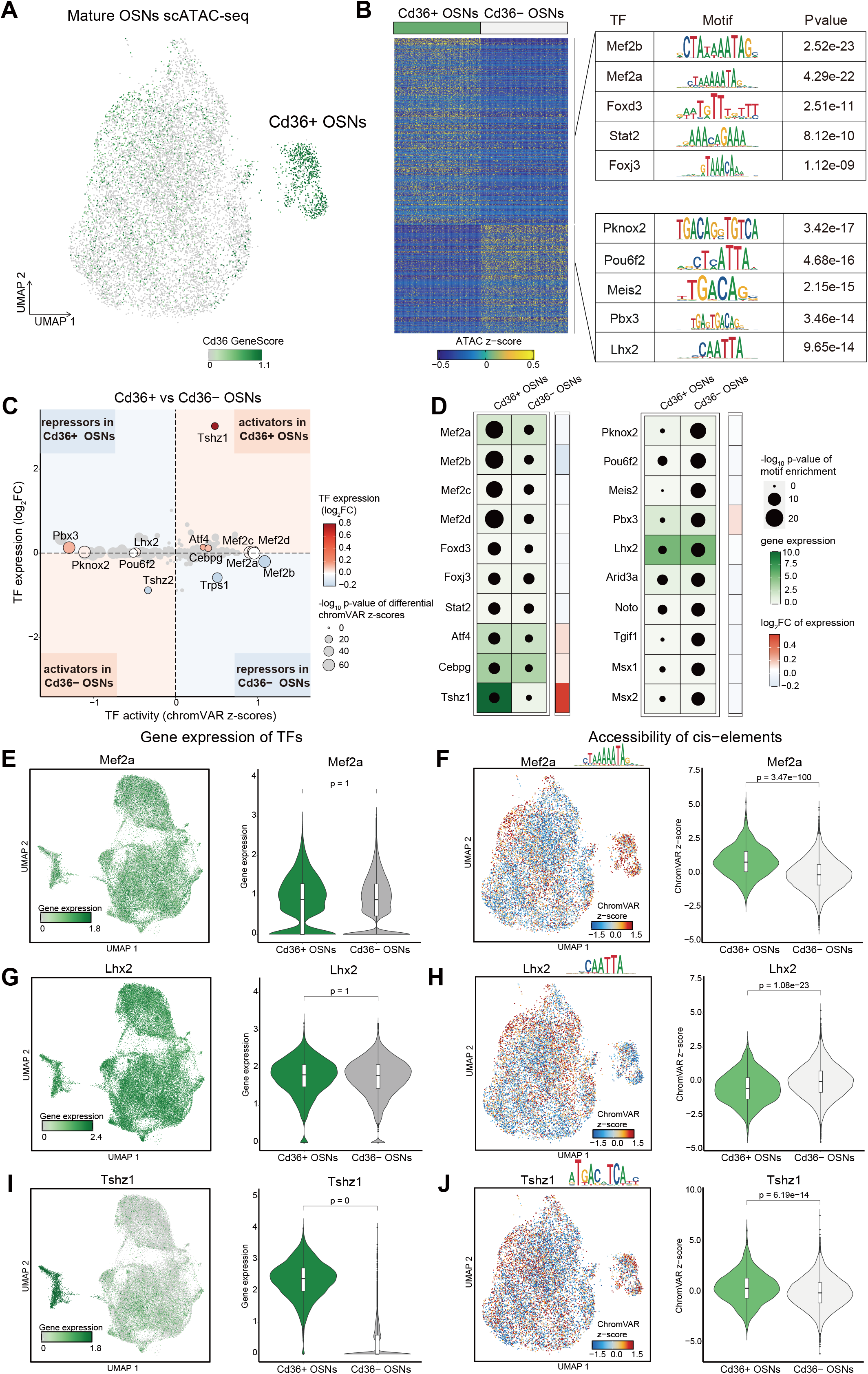
Differential chromatin accessibility in Cd36+ OSNs. (A) UMAP plot showing the subclustering result of the scATAC-seq data from mOSNs. Each cell is colored by the gene activity score of Cd36. The cluster with high accessibility around Cd36 is annotated as Cd36+ OSNs. (B) Left: heatmap of differentially accessible peaks between Cd36+ OSNs and Cd36- OSNs (computed by a logistic regression model in Signac package, Bonferroni adjusted p-value < 0.05). Shown are z-transformed chromatin accessibility of differentially accessible peaks in 500 cells sampling from Cd36+ OSNs and Cd36- OSNs. Right: enriched motifs of upregulated peaks in Cd36+ OSNs and Cd36- OSNs, respectively. (C) Scatter plot showing the relationship between Cd36+ vs Cd36- OSNs differential expression levels (log2FoldChange) and differential chromVAR activities for enriched TFs. The dot color represents the log2 fold change of gene expression and the dot size represents the -log10 (p-value) obtained from differential TF chromVAR activity analysis. (D) Dot plots showing Top-10 enriched TF motifs in up-regulated peaks of Cd36+ OSNs (Left) and Cd36- OSNs (Right), respectively. The dot size represents -log10 (p-value) obtained from motif enrichment analysis in up-regulated peaks. The gene expression levels of TFs in Cd36+ OSNs and Cd36- OSNs were shown from light green to dark green. The log2 fold change of TFs expression between Cd36+ and Cd36- OSNs was shown beside each dot plot. (E) UMAP plot and violin plot showing the gene expression of Mef2a. (F) UMAP plot and violin plot showing the Mef2a motif activity measured by chromVAR z-scores. (G) UMAP plot and violin plot showing the gene expression of Lhx2. (H) UMAP plot and violin plot showing the Lhx2 motif activity measured by chromVAR z-scores. (I) UMAP plot and violin plot showing the gene expression of Tshz1. (J) UMAP plot and violin plot showing the Tshz1 motif activity measured by chromVAR z-scores.

Focusing on the MEF2 family with the most significant motif enrichment, we identified Mef2a with the highest expression level compared to Mef2b, Mef2c, and Mef2d (**Figures S6A-S6D**). However, the expression levels of Mef2 genes did not significantly differ between Cd36+ and Cd36- OSNs. On the contrary, noticeable differences in chromatin accessibility were observed at peaks containing MEF2 binding motifs, as quantified by chromVAR z-scores, between Cd36+ and Cd36- OSNs (**Figures S6A-S6D; Figures 3E and 3F**). The transcription factors that are enriched in the down-regulation peaks showed the same pattern, such as Lhx2, the one with consistently high expression levels both in Cd36+ and Cd36- OSNs (**Figure 3D**; **Figures 3G and 3H**). It has been shown that Lhx2 plays a crucial role in the selection of ORs by binding at OR enhancers, the ‘Greek island’, and mediating trans interactions between OR promoters to contribute to the formation of cell-specific chromatin compartments (Hirota et al., 2007; Monahan et al., 2019; Monahan et al., 2017). While these transcription factors did not exhibit apparent differences in their transcription levels between Cd36+ and Cd36- OSNs, they displayed significant variations in chromatin accessibility at their motif binding *cis*-elements. In contrast to Mef2a, we identified that Tshz1 showed high expression specificity in Cd36+ OSNs, and the Tshz1 motif exhibited significantly higher ChromVAR z-scores in Cd36+ OSNs compared to Cd36- OSNs (**Figures 3I and 3J**). In summary, through a comprehensive analysis of epigenome profiles, we have unveiled the unique chromatin landscape of Cd36+ OSNs, and their corresponding trans-factors that may govern cellular diversity. Interestingly, these results suggest that regulatory differences driven by most of the trans-factors may not depend on their expression specificity, and that the regulatory mechanisms of OSNs diversity remain to be elucidated.

### Trajectory analysis reveals the temporal expression of trans-factors during OSN development

Having established the co-expression patterns of Cd36 and a specific subset of ORs, as well as the unique *cis* and *trans* signatures of Cd36+ OSNs, we proceeded to investigate their coordinated regulation during the development and maturation of OSNs. In addition, we wonder whether the transcription factors that showing divergent binding activities in the mature OSNs stage had expression differences at the previous development stage, which may explain the controversy.

To answer these questions and elucidate the timing of the initiation of Cd36+ OSN diversity, we performed trajectory analysis on cells of the olfactory neuronal lineage (**Methods and Figures S7A and 7B**). We identified a developmental period prior to OSN maturation, during which OSNs began to express modest amounts of olfactory marker protein (Omp) but showed lower expression of adenylate cyclase 3 (Adcy3), indicating that they are a transitional stage of mature OSNs (Lyons et al., 2013) (**Figure S7C**). These OSNs were defined as Adcy3- OSNs (**Figure S7D**). RNA velocity analysis revealed the developmental trajectory of the olfactory neuronal lineage from globose basal cells (GBCs) to mature OSNs (**Figure 4A**). Furthermore, at the Adcy3- OSNs stage, Cd36+ OSNs were clearly distinguished from Cd36- OSNs (**Figure 4A; Figure S7E**), and Tshz1 was also significantly expressed in the precursor cells of Cd36+ OSNs at the Adcy3- OSNs stage (**Figure 4B**). Adcy3- OSNs with high Tshz1 expression levels were classified as precursor cells of Cd36+ OSNs, while the remaining cells with low or no Tshz1 expression were assigned as precursor cells of Cd36- OSN (**Methods and Figure S7F**). Consequently, we obtained the development trajectories of Cd36+ and Cd36- OSNs, respectively (**Figure 4C**).

**Figure 4.**
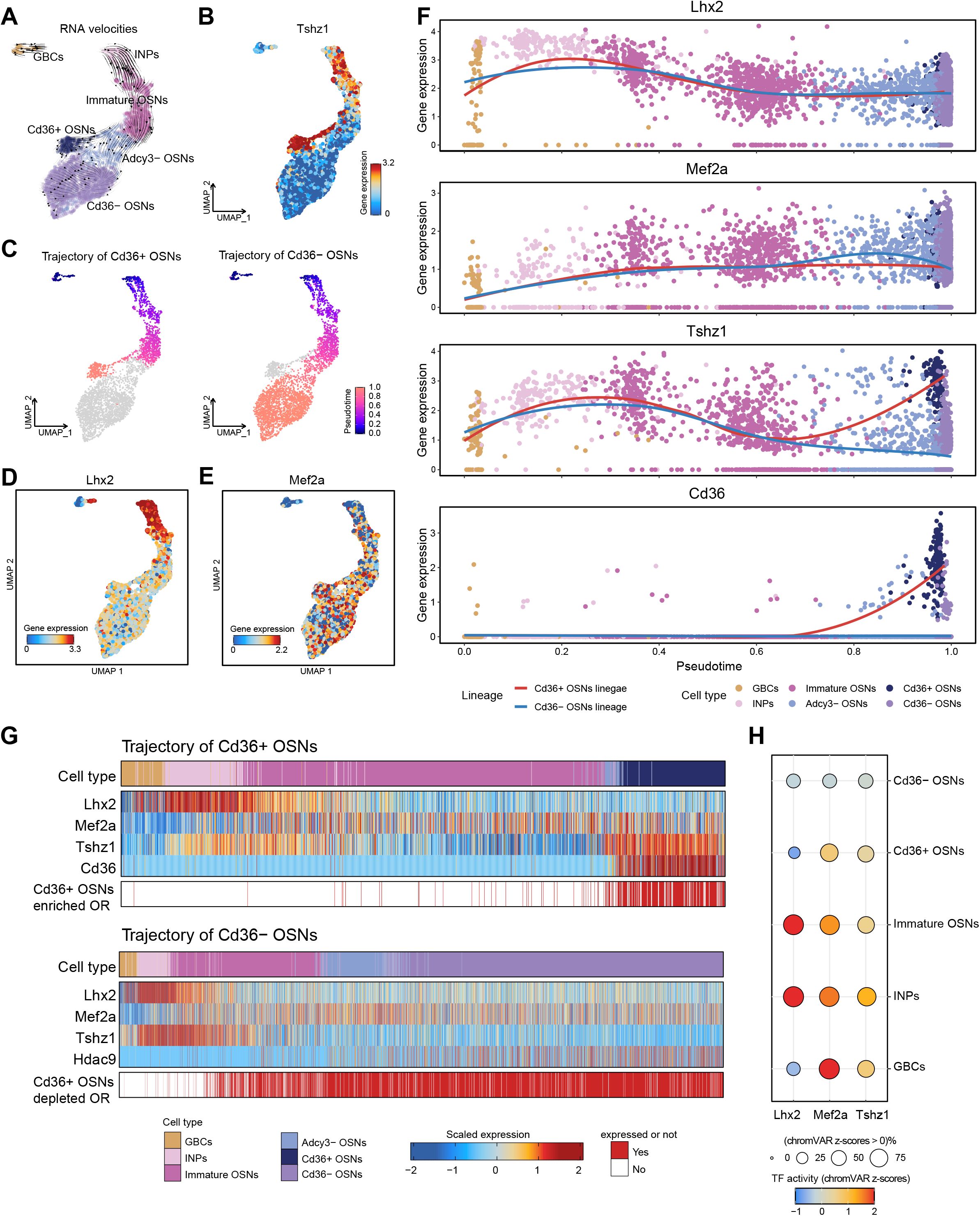
The expression patterns of important transcription factors during OSNs development. (A) RNA velocities, estimated by scVelo, projected into a UMAP embedding of the olfactory neuronal lineage (HBCs are excluded). (B) UMAP plot showing the gene expression of Tshz1. (C) UMAP plots for the trajectory of Cd36+ OSNs (left) and Cd36- OSNs (right), respectively. Cells in a specific trajectory are colored by pseudotime derived from RNA velocity analysis, while cells that are not in the given trajectory are colored gray. (D) UMAP plot showing the gene expression of Lhx2. (E) UMAP plot showing the gene expression of Mef2a. (F) Expression of Lhx2, Mef2a, Tshz1 and Cd36 along the pseudotime of Cd36+ OSNs and Cd36- OSNs trajectory, respectively. (G) Top: Expression of Lhx2, Mef2a, Tshz1, Cd36 and Cd36+ OSN-enriched OR genes along the pseudotime of Cd36+ OSNs trajectory. Cells in the trajectory of Cd36+ OSNs are sorted by pseudotime in ascending order. A Cd36+ OSN-enriched OR gene is considered expressed if at least 4 UMIs are detected. Bottom: Expression of Lhx2, Mef2a, Tshz1, Hdac9 and Cd36+ OSN-depleted OR genes along the pseudotime of Cd36- OSNs trajectory. Cells in the trajectory of Cd36- OSNs are sorted by pseudotime in ascending order. A Cd36+ OSN-depleted OR gene is considered expressed if at least 4 UMIs are detected. (H) Dot plot showing Lhx2, Mef2a, and Tshz1 motif activity in each olfactory neuron lineage-related cell type, demonstrated by average chromVAR z-scores. The dot size represents the proportion of cells in that cell type that have a chromVAR z-score greater than 0. The dot color represents the average chromVAR z-score in that cell type.

Along the developmental trajectories, we investigated the expression levels of transcription factors showing significant motif enrichment in differential accessibility regions between Cd36+ and Cd36- OSNs. Interestingly, we found that Lhx2 is the earliest expressed regulator, which is expressed at the GBCs stage and highest expressed at the immediate neuronal precursors (INPs) stage (**Figures 4D and 4F**). However, there is no significant difference in the expression level of Lhx2 between the two lineages along the developmental trajectory. Although significant differential accessibility at Mef2a binding sites was observed, Mef2a started to express at the INPs stage and increased along the development trajectory, but neither shows any expression difference between the two lineages (**Figures 4E and 4F**). In contrast to Lhx2 and Mef2a, the expression levels of Tshz1 increased in the early developmental stage of INPs and subsequently declined for most cells in the immature OSNs stage (**Figures 4B, 4F and 4G**). Tshz1 remained constantly expressed only in Cd36+ OSN precursors at the Adcy3- OSNs stages and eventually became specifically expressed in Cd36+ OSNs, demonstrating that the expression divergence of Tshz1 was initiated in the immature stage, when OR expression was determined (**Figures 4B, 4F and 4G**).

In combination with the binding activities of these three TFs, we found that both binding activities and expression levels tend to be higher in the early developmental stage (e.g., INP stage) (**Figures 4G and 4H**). Gradually reaching the mature OSNs stage, Mef2a and Tshz1 are more active in Cd36+ OSNs and Lhx2 is more active in Cd36- OSNs (**Figure 4H**). These results showed that temporal differential expression of trans-factors during OSN development could not explain their binding activity differences, suggesting extraordinary regulatory mechanisms driving the transcription factor binding divergence in OSN populations.

### Key regulators of cell-fate determination in Cd36+ OSNs and Cd36- OSNs

Intriguingly, both Lhx2 and Mef2a exhibited continuous expression since the immature OSNs stage and did not show obvious expression differences between Cd36+ and Cd36- OSNs (**Figures 4D-4F**). How trans-factors with homogeneous expression play divergent roles in the OSN population, is of great interest.

In the transcriptome analysis, we found that Hdac9, which is enriched in deacetylase activity GO terms, was significantly higher expressed in Cd36- OSNs (**Figure 1B**). Interestingly, along the trajectory analysis, we found that Hdac9 was robustly expressed in a lineage-specific manner at the mature stage of Cd36- OSNs (**Figure 5A**). By immunofluorescence analysis, HDAC9 protein was indeed significantly higher in Cd36- OSNs compared with Cd36+ OSNs (**Figures 5B and 5C**). Furthermore, we found a Cd36- OSNs specific peak on the intron region of Hdac9, located 86, 750 bp downstream from the Hdac9 promoter, and overlapping with the Lhx2 ChIP-seq peak signal (**Figure 5D**). Peak-gene linkage analysis also showed that this enhancer peak has the strongest linkage with Hdac9 expression (**Figure 5D**). These results indicate that Lhx2 may promote the expression of Hdac9 in Cd36- OSNs. To confirm this regulatory relationship, we analyzed bulk RNA-seq datasets from Lhx2 knockout mice (Monahan et al., 2017), and revealed a significant downregulation of Hdac9 expression in mature OSNs in the absence of Lhx2 (**Figure 5E**).

**Figure 5.**
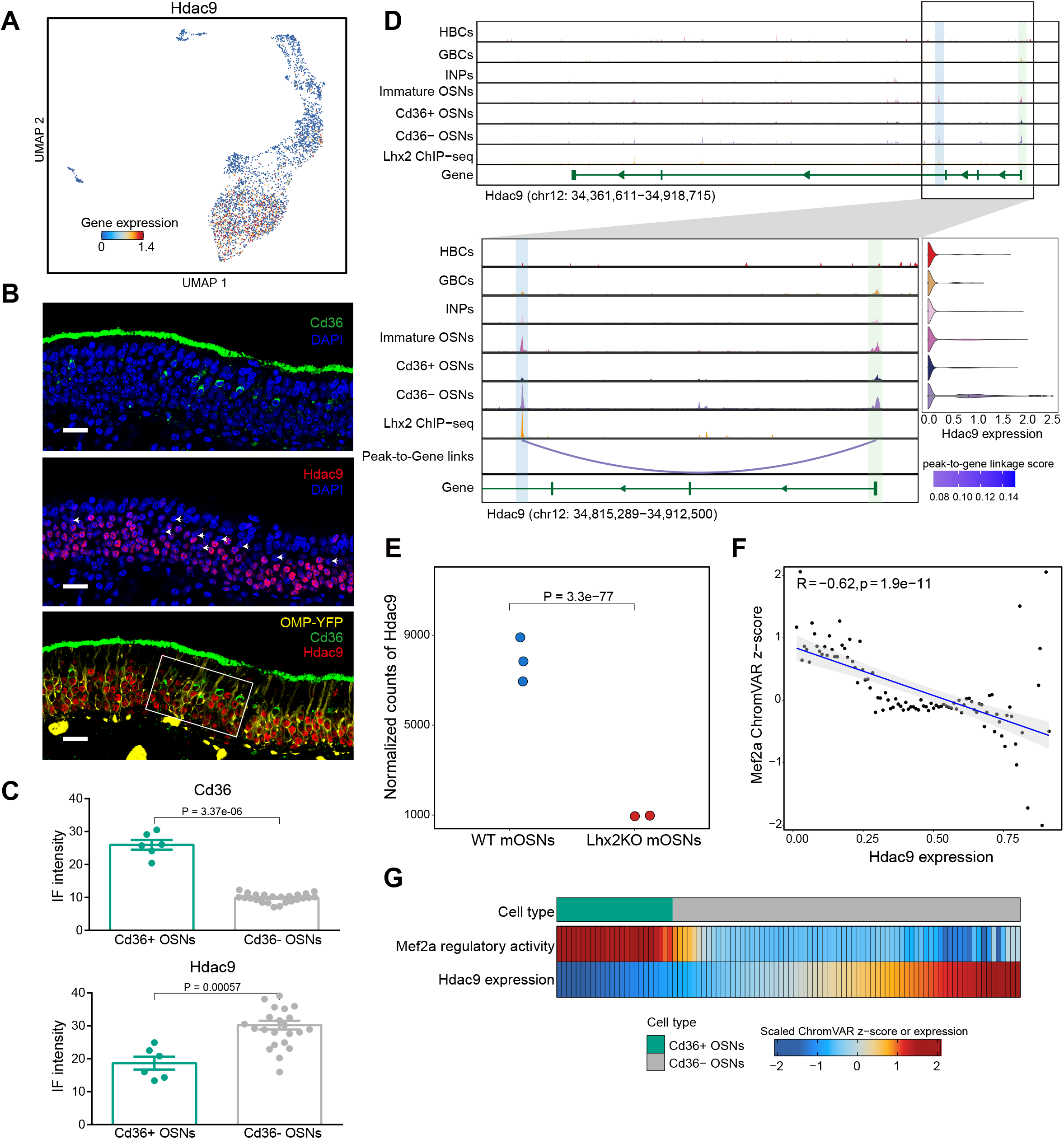
Key regulators of fate determination in Cd36+ OSNs and Cd36- OSNs. (A) UMAP plot showing the expression of Hdac9. (B) Confocal images showing CD36 (green) and HDAC9 (red) immunostaining of a mouse olfactory epithelial section. DAPI indicates nuclear staining (blue). The fluorescence signal of the OMP protein is preserved (yellow). Scale bar, 20 μm. The white box applies to Figure 5C. (C) Bar plot showing the immunofluorescence intensity for CD36 and HDAC9, quantified using the mean gray value. Each green dot represents a Cd36+ OSN and each gray dot represents a Cd36- OSN. Significance was calculated by the Wilcoxon Rank Sum test. Data represent mean ±SEM. (D) Genome tracks showing the peak with Lhx2 ChIP-seq signal is positively linked to Hdac9 expression. Tracks from top to bottom are normalized chromatin accessibility around the Hdac9 gene in HBCs, GBCs, INPs, immature OSNs, Cd36+ OSNs and Cd36- OSNs, Lhx2 binding signals profiled by ChIP-seq obtained from Monahan (Monahan et al., 2017), gene annotation, and peak-to-gene links. The promoter region of Hdac9 with high chromatin accessibility in Cd36- OSNs is highlighted in light green, while the enhancer region is highlighted in blue. Beside the genome track is the violin plot showing Hdac9 expression in each cell type of olfactory neuronal lineage. (E) Normalized gene expression of Hdac9 in wild-type and Lhx2 KO mOSNs obtained from Monahan et al (Monahan et al., 2017). (F) Scatter plot showing the relationship between the Mef2a motif activity (ChromVAR z-score) and Hdac9 expression. (G) Smoothed heatmap showing the relationship between the Mef2a regulatory activity and Hdac9 expression mature OSNs.

Hdac9, as a class II histone deacetylase, is specifically expressed in Cd36- OSN, implying its potential role in epigenetically repressing genes that contribute to Cd36+ OSN diversity. Besides, previous studies have shown that Hdac9 can interact with Mef2a and function as a transcriptional corepressor, to suppress the activity of Mef2a and reduce the expression of its target genes (Haberland et al., 2007; Mejat et al., 2005; Minisini et al., 2022; Zhang et al., 2001). The lineage-specific expression of Hdac9 may explain the regulatory and binding diversity of Mef2a. In conjunction with Hdac9 expression, we found a negative correlation between the TF activity of Mef2a and Hdac9 expression, indicating that the elevated TF activity of Mef2a in Cd36+ OSNs may result from a lack of Hdac9-mediated repression (**Figures 5F and 5G**).

In summary, we found the binding activity divergence of Mef2a may be driven by the lineage-specific expression of Hdac9 in Cd36- OSNs, where the regulatory activity of Mef2a was abrogate on Cd36+ OSN up-regulated genes.

### Regulation networks and key regulators in promoting lineage-specific expression of Cd36

We further built a gene regulatory network (GRN) for Cd36+ OSNs (**Method; Figure 6A**), which identified Mef2a and Tshz1 as the key regulators for Cd36+ OSN up-regulated genes. Interestingly, we found Tshz1 is also regulated by Mef2a via cis-element binding (**Figure 6A**). When we examined the chromatin accessibility profiles around the Tshz1 gene, we found a Cd36+ OSN-specific enhancer region containing the Mef2a binding motif and exhibiting the strongest linkage with Tshz1 expression (**Figure 6B**).

**Figure 6.**
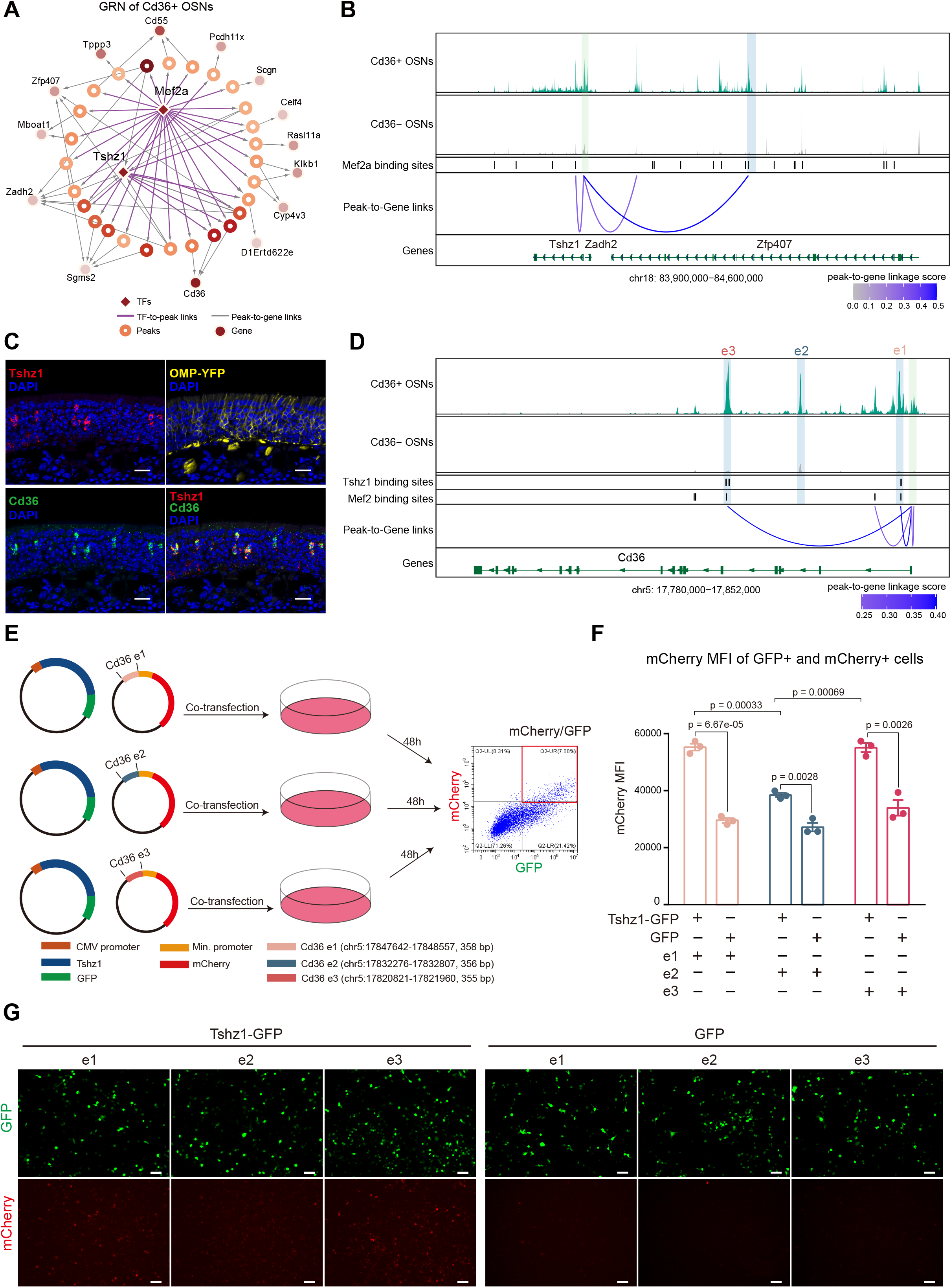
Cellular diversity of Cd36+ OSNs is contributed by the transcription factors Mef2a and Tshz1. (A) Putative gene regulatory network for Cd36+ OSNs with key TFs in the center. Up-regulated peaks in Cd36+ OSNs are shown by hollow circles, and the color indicates the differential accessibility of the peak between Cd36+ OSNs and Cd36- OSNs. Up- regulated genes in Cd36+ OSNs are shown in solid circles, and the color indicates the differential expression of the gene between Cd36+ OSNs and Cd36- OSNs. TF-to-peak links are based on motif searching, and peak-to-gene links are based on the correlation of peak accessibility and gene expression. (B) Genome tracks showing peaks with Mef2a motifs are positively linked to Tshz1 expression. Tracks from top to bottom are normalized chromatin accessibility around the Tshz1 gene in Cd36+ OSNs and Cd36- OSNs, Mef2 binding sites, peak-to-gene links (z score >= 0.27 are shown) and gene annotations. The promoter region of the Tshz1 gene with high chromatin accessibility in Cd36+ OSNs is highlighted in light green, while enhancer regions are highlighted in blue. (C) Confocal images of mouse main olfactory epithelium sections showing the colocalization of Tshz1 (red) and Cd36 (green) in mature OSNs using multiplexed fluorescent RNAscope probes. DAPI indicates nuclear staining (blue). The fluorescence signals of the Omp protein are kept (yellow). Scale bar, 20 μm. (D) Genome tracks showing peaks with Tshz1 and Mef2 motifs are positively related to Cd36 expression. Tracks from top to bottom are normalized chromatin accessibility around the Cd36 gene (chr10: 17780000-17852000) in Cd36+ OSNs and Cd36- OSNs, Tshz1 binding sites, Mef2 binding sites, peak-to-gene links (z score >= 0.2 are shown) and gene annotation. The promoter region of the Cd36 gene with high chromatin accessibility in Cd36+ OSNs is highlighted in light green, while enhancer regions are highlighted in blue. (E) Experimental overview of the in vitro CREs-reporter assay. N2a cells were co-transfected with two plasmids constructed using full-length Tshz1-GFP (control GFP) and Cd36 enhancer (1-3)-mCherry. After 48 hours of transfection, cells were visualized using fluorescence microscopy and flow cytometry. (F) Bar plots showing the mCherry mean fluorescence intensity (MFI) in the fluorescence-activated sorted GFP+/mCherry+ cells for different experimental conditions. Significance was calculated by unpaired Student’s t-test. Data represent mean ±SEM. (G) Co-transfection experiments were visualized by fluorescence microscopy. Scale bar, 50 μm.

Tshz1 demonstrated remarkable expression specificity in Cd36+ OSNs compared to other *trans*-regulators. To validate the specific expression of Tshz1 in Cd36+ OSNs, we performed RNAscope *in situ* hybridization and observed effective colocalization of Tshz1 fluorescence signals (in red) with Cd36 in YFP+ OSNs (**Figure 6C**). As expected, the fluorescence signal of Tshz1 was lower in YFP-progenitor cells and Cd36 was abundantly expressed on the surface of the olfactory epithelium and the cilia of OSNs, consistent with previous observations (Lee et al., 2015) through immunofluorescence staining. Approximately 5% of YFP+ OSNs exhibited Cd36 signals (**Figure 6C**), confirming the proportion of Cd36+ OSNs determined by scRNA-seq and scATAC-seq.

The GRN analysis identified that Tshz1 directly regulated the marker gene Cd36. In the trajectory analysis, we also observed that Cd36 expression occurred after the expression of Tshz1 (**Figure 4G**). We further identified three *cis*-regulatory elements (e1, e2 and e3) located within the gene body of Cd36. Notably, e1 and e3, which contain Tshz1 binding motifs, exhibited strong linkage with Cd36 expression (**Figure 6D**). To confirm this regulatory relationship between Tshz1 and Cd36, we conducted a *cis*-regulatory elements (CREs)-reporter assay in vitro to determine if Tshz1 binds to CREs containing Tshz1 motifs and drives Cd36 expression (**Methods; Figure 6E**). Briefly, we constructed two types of plasmids. One enabled the overexpression of Tshz1 with its expression monitored by GFP, and the other contained reporter plasmids with one of the three potential Cd36 enhancer sequences inserted upstream of the minimal promoter of the reporter gene (mCherry). Subsequently, we co-transfected the overexpression- and reporter-plasmids into the Neuro2a (N2a) cell line. After 48 hours, the expression of the reporter gene mCherry in different cells was detected by flow cytometry. As expected, when Tshz1 was overexpressed, the mean fluorescence intensity (MFI) of mCherry was significantly increased in Tshz1-GFP+/mCherry+ dual-plasmid co-transfected cells, compared to those transfected with empty plasmid (GFP) and reporter plasmids (**Figure 6F; Figures S8A and S8B**). Furthermore, when Tshz1 was overexpressed, e1 and e3 containing the Tshz1 motif exhibited significantly higher expression levels of mCherry reporter compared to that of e2 without the Tshz1 motif (**Figures 6F and 6G**). These findings support the notion that Tshz1 is capable of driving Cd36 expression through its interactions with specific CREs. We have identified novel CREs in OSNs that are responsible for mediating the regulatory effect of Tshz1 on Cd36 expression. Tshz1 plays a crucial role in promoting Cd36 expression and driving the cellular diversity of OSNs. Collectively, we systematically revealed the regulatory relationship, composited with transcription factors, cis-elements and their corresponding target genes, in the regulation of cellular diversity of OSNs.

## Discussion

In this study, we conducted a comprehensive analysis of the transcriptome diversity and spatial distribution of Cd36+ OSNs in the mouse olfactory epithelium. The unique molecular features of Cd36+ OSNs indicate their programmed cellular diversification, which contribute to their functional diversity in olfaction. By integrating single-cell transcriptome and epigenome profiles, we further identified both *cis* and *trans*-regulatory signatures in Cd36+ OSNs. Intriguingly, we found that the majority of the transcription factors revealed by the motif enrichment analysis were not specific or highly expressed in the Cd36+ OSN population. This finding contrasts with the cell fate determination during development, where lineage-specific transcription factors are typically expressed in a lineage-specific manner. Our results suggest that the regulation of cellular diversity may involve different principles, such as post-translational modification and the combinatorial action of transcription factors.

Previous studies showed that cyclic adenosine monophosphate (cAMP) acts as a key second messenger in signal transduction pathways, which is essential for axon guidance and neuronal plasticity in OSNs (Imai et al., 2006; Nakashima et al., 2021; Nakashima et al., 2013; Takeuchi and Sakano, 2014). It has been shown that cAMP signaling can modulate the interaction between HDAC (class II histone deacetylases) and MEF2 transcription factors, which play a critical role in regulating gene expression targeted by MEF2 (Belfield et al., 2006; Du et al., 2008; He et al., 2020; Jebessa et al., 2019). In the transcriptome comparison, we found the expression of Neuropilin1 (Nrp1) is lower in Cd36+ OSNs, indicating lower basal cAMP level in Cd36+ OSNs (**Figure S9A**) (Col et al., 2007; Imai et al., 2006; Nakashima et al., 2021; Nakashima et al., 2013). Furthermore, we also found that adenylyl cyclase type III (Adcy3), an enzyme for cAMP synthesis, exhibits expression differences between Cd36+ and Cd36-OSNs (**Figure S9B**). These results indicate that the cAMP level differences in OSNs may also contribute to the interaction between Hdac9 and Mef2a, which may further contribute to the transcriptome and epigenome diversity of Cd36+ and Cd36-OSNs. The signaling-dependent modification and activation of trans-factors may resolve the limitation on cellular diversity imposed by the number of encoded transcription factors, requiring further investigations. Our findings shed light on the mechanisms driving cellular diversity in the mouse OSNs and provide new insight into the complex regulatory processes underlying complex organ development and function.

Besides the question on how cellular diversity is regulated, another intriguing question is how cellular diversity evolved. We further investigated the expression patterns of Cd36 and Tshz1 in olfactory mucosa across multiple species, including mice, rats, dogs, marmosets and humans (Saraiva et al., 2019). We observed a strong correlation in the expression levels of Cd36 and Tshz1 across these species. Specifically, rodents exhibited the highest expression levels, canines showed moderate levels, and primates displayed the lowest expression levels (**Figure S9C**). These findings provide independent evidence supporting the role of Tshz1 in regulating the expression of Cd36. Furthermore, the species-specific expression patterns of Cd36 and Tshz1 imply that the function of Cd36+ OSNs may be specific to rodents, highlighting potential evolutionary differences in olfactory systems among species. The absence or reduced expression of Cd36 and Tshz1 in the olfactory epithelium in other species (such as dogs, marmosets and humans) suggests that Cd36+ OSNs may be absent or less abundant compared to mice. The two scenarios could not be distinguished with bulk RNA-seq data alone. We examined scRNA datasets from human OE (Durante et al., 2020; Finlay et al., 2022) and found no evidence of OSNs with Cd36 expression (data not shown). However, due to the limited sample area of the human olfactory epithelium, we cannot conclusively state that Cd36+ OSNs are entirely absent in humans. Large-scale single-cell profiling studies in other species are necessary to investigate the conservation of Cd36+ OSN population, and comparative analysis across multiple species will provide further insights into the evolution of cellular diversity and its relationship to species-specific adaptations.

## Availability of data and materials

Sequencing data is available on the Gene Expression Omnibus repository under accession number GSE224604.

Link for GSE224604 is https://www.ncbi.nlm.nih.gov/geo/query/acc.cgi?acc=GSE224604 (reviewer token: ——). The Code used for data analysis and visualization is available at https://github.com/PeggySze/Cd36_OSNs.

## Acknowledgments

We thank Rui Zhang (Sun Yat-sen University), Xiaowei Zhu (City University of Hong Kong), and Ningyi Shao (University of Macau) for constructive comments and suggestions on the manuscript, as well as the members of our lab for their helpful advice.

## Author contributions

J.X., J.Y. and P.S. designed and led the project. D.P., J.Y., Y.X. and A.L. performed mouse breeding and tissue collection. J.Y., Y.L., Y.Z., T.X. and W.Z. performed the molecular experiments and interpreted the data. P.S., J.Y., Z.A. and Z.T. analyzed and interpreted the data. J.Y., P.S. and J.X. drafted and edited the manuscript, and all the other authors commented on the manuscript.

## Online Methods

### Mice

OMP-ChR2-YFP (OCY58) mice were obtained from Li laboratory (Xuzhou Medical University). PWK/PhJ mice were obtained from Jackson Laboratory with stock number 003715. F1 hybrid mice were produced by crossing OMP-ChR2-YFP (maternal strain) and PWK/PhJ (paternal strain) mice. The F1 hybrid mice were used for the single-cell and RNAscope experiments. All mice were housed in barrier facilities in a 12-hour light/dark cycle with free access to standard mouse diet and water. All experiments involving animals were performed following guidelines approved by the Institutional Animal Care and Use Committee of Sun Yat-Sen University, P.R. China.

### Dissection of mouse olfactory epithelium and isolation of mOSNs

F1 hybrid mice (6-8 mice with either sex in 19 days or 8 weeks for each single-cell library) were deeply anesthetized and perfused transcardially with 20 ml of ice-cold phosphate-buffered saline (PBS) (Oberland and Neuhaus, 2014). Mice were then rapidly decapitated, and turbinate tissues were washed and immersed in ice-cold PBS. Turbinate tissues were minced into small pieces and disaggregated in 0.25% Trypsin-EDTA (1x) (GIBCO) with gentle agitation at 37 °C for 20 min. After adequate disaggregation, equal volume DMEM containing 10% fetal bovine serum (FBS, GIBCO) was added to stop the disaggregation reaction, and then the dissociated cells were filtered through a 40 μm cell strainer (ThermoFisher). After centrifugation at 300 g for 5 min, dissociated cells were resuspended in ice-cold PBS and centrifuged at 300 g at 4 °C for 5 min. The cell pellets were then resuspended in 1×Red Blood Cell Lysis Solution (Miltenyi Biotec) and equal volume ice-cold PBS was added and centrifuged at 300 g at 4 °C for 5 min. Dead Cell Removal Kit (Miltenyi Biotec) was applied to exclude dead cells. To enrich olfactory sensory neurons, YFP+ cells (mOSN) were sorted by MoFlo Astrios EQ cell sorter (Beckman Coulter, USA).

### Single-cell RNA-seq library preparation and sequencing

Single-cell sequencing libraries were prepared using the Chromium Single Cell 3′ Reagent Kit V3.1 (10x Genomics) according to the manufacturer’s guidelines. Sequencing library fragments were examined using the Agilent High Sensitivity DNA Kit (Agilent) and quantified via qPCR by the KAPA library quantification kit (Roche). All single-cell libraries were sequenced on a Novaseq 6000 system (Illumina, US) (minimum read lengths: Read1 = 28 cycle, Index (i7) = 8 cycle, Index (i5) = 0 cycle, Read2 = 91 cycle for v3.1 kits).

### Single-cell ATAC-seq library preparation and sequencing

Sinlge-cell ATAC-seq libraries were prepared using the Chromium Single Cell ATAC Reagent Kit V1.1 (10x Genomics) according to the manufacturer’s guidelines. Sequencing library fragments were examined using the Agilent High Sensitivity DNA Kit (Agilent) and quantified by qPCR with the KAPA library quantification kit (Roche). All libraries were sequenced on a Novaseq 6000 system (Illumina, US) (minimum read lengths: Read 1 = 50 cycle, Index (i7) = 8 cycle, Index (i5) = 16 cycle, Read 2 = 50 cycle).

### RNAscope in situ hybridization

8-week-old mice were sacrificed and turbinates were immediately fixed in freshly prepared 4% PFA 24h at 4 °C. Turbinate tissue was immersed in 0.45M EDTA for 12 h, 20% sucrose for 6 h and 30% sucrose for 6 h at 4 °C. Turbinate was embedded in OCT, frozen and stored at -80 °C until sectioning. The turbinate was cut on a cryostat microtome (Leica CM 1950, Leica Biosystems) in 10 μm coronal sections. RNAscope staining was performed according to the manufacturer’s protocol (RNAscope® Multiplex Fluorescent Reagent Kit v2, ref. 323100, Advanced Cell Diagnostics). Pretreatment was carried out according to guidelines for fixed-frozen tissue and included postfixation, dehydration, hydrogen peroxide treatment and 20 min Protease III treatment. For the F1 hybrid mice, the endogenous OMP-YFP protein signal was maintained because of the lack of target retrieval. The sections were labelled with probes for Tshz1 and Cd36 (RNAscope® Probe-Mm-Tshz1, ref. 494291, RNAscope® Probe-Mm-Cd36-C2, ref. 464431-C2, both from Advanced Cell Diagnostics). Probes were visualized with Opal fluorophores (Opal 570 Reagent, ref. ASOP570, Opal 690 Reagent, ref. ASOP690). Sections were counterstained with DAPI and mounted with ProLongTM Gold antifade (ref. P36935, Invitrogen). Slides were imaged with a Zeiss LSM880 confocal microscope equipped with 405, 514, and 633 nm laser lines, using a 40 ×1.0 NA oil-immersion objective.

### Cis-regulatory elements (CREs)-reporter assay

Reporter gene expression protocol as previously described with a few modifications (Melnikov et al., 2014; Nakashima et al., 2020; Noack et al., 2022). Neuro2a cells were cultured at a density of 2.0 ×10^5^/well in 6-well plates transiently transfected with 1µg of plasmid/well, which included 500 ng of CREs-min. promoter-mCherry (reporter) plus 500 ng of the pCMV-TSHZ1-GFP expression plasmids or the reporter plus pCMV-GFP (as control), using the Lipofectamine 3000 (Invitrogen) transfection reagent according to the manufacturer’s protocol. 48 h after transfection, cells were visualized by fluorescence microscopy and applied to the FACS analysis. Experiments were performed 3 times on independent N2a cultures.

### scRNA-seq data processing and cell type identification

For each scRNA-seq dataset, raw reads were aligned to the *Mus musculus* reference genome mm10 (Ensembl 98) and converted into gene-barcode matrix using 10x Genomics Cell Ranger (Zheng et al., 2017) version 4.0.0. Count data were further processed using the R package Seurat (Hao et al., 2021) version 4.1.0. High-quality cells that expressed 400-5,000 genes and had a mitochondrial content <10% were retained for downstream analysis. Single-cell RNA-seq datasets across different replicates and time points were integrated using the standard integration workflow (https://satijalab.org/seurat/articles/integration_introduction.html). Briefly, count data from each sample were log-normalized using the NormalizeData() function, and the top 3000 variable features were identified using the FindVariableFeatures() function. To integrate scRNA-seq datasets, a set of anchors was identified using the FindIntegrationAnchors() function with 3000 genes. Following the integration of the four scRNA-seq datasets using these anchors, the data were scaled using the ScaleData() function. The principal component analysis was then performed on the integrated assay. The top 40 principal components (PCs) were selected based on the elbow plot and further used for clustering at a resolution of 0.8 and the Uniform Manifold Approximation and Projection (UMAP) dimensional reduction was employed for visualization. This preliminary clustering yielded 31 clusters. Cluster-specific gene markers were identified using the Wilcoxon rank sum test implemented in the FindAllMarkers() function. We first annotated cells that do not belong to the olfactory neuronal lineage using the following marker genes: Sustentacular cells (*Sox2*, *Ermn*); Ensheathing glia (*Atp1a2*, *Fabp7*); Microvillar cells (*Ascl3*, *Cftr*); Bowman’s gland (*Sox9*, *Sox10*); Periglomerular cells (*Pebp1*, *Calb2*); Pericytes (*Eng*, *Sox17*); Brush cells (*Krt18*, *Trpm5*); Osteogenic cells (*Col1a1*, *Bglap*); B cells (*Cd37*, *Cd79a*); Late activated neural stem cells (*Hmgb2*, *Top2a*); Erythrocytes (*Hbb-bs*, *Hbb-bt*); Basophils (*Mcpt8*); Neutrophils (*S100a9*, *S100a8*); Monocytes (*Lyz2*, *S100a4*); Macrophages (*C1qa*, *Ms4a7*). Due to the incapability to distinguish between GBCs and INPs based on the preliminary clustering, 10 cell clusters that corresponded to cells in the olfactory neuronal lineage (HBCs, GBCs, INPs, Immature OSNs, and mature OSNs) were extracted for further subclustering analysis. The standard integration workflow of Seurat was applied to the olfactory neuronal lineage subclustering analysis as mentioned above with 2000 anchor features and the top 35 principal components for clustering at a resolution of 0.8. Cells in the olfactory neuronal lineage were annotated using the following marker genes: HBCs (*Cebpd*, *Krt5*); GBCs (*Ascl1*, *Kit*); INPs (*Neurod1*, *Sox11*); Immature OSNs (*Gng8*, *Gap43*); Mature OSNs (*Omp*, *Syt1*).

### Subclustering analysis of mature OSNs scRNA-seq data

Based on the annotations of the integrated scRNA-seq data, mature OSNs were extracted for further analysis. We first identified the OR gene expressed in each mature OSN. Using the expert curation of mouse olfactory receptor gene repertoires from Barnes IHA et al (Barnes et al., 2020), we annotated 1140 OR genes in the *Mus musculus* reference genome mm10 (Ensembl 98). Of these 1140 OR genes, an OR gene was considered expressed in any given mature OSN if at least 4 UMIs were detected. Mature OSNs expressing a single OR gene were retained for further downstream analysis. Next, the standard integration workflow of Seurat was applied to the subclustering analysis of retained mature OSNs with 1500 anchor features (exclude mitochondrial genes, OR genes, highly-expressed lncRNA Malat1, and sex-specific gene Xist) and the top 40 principal components for clustering at a resolution of 0.2. It yielded 6 clusters, 2 of which had a relatively low number of expressed genes and were excluded. Retained mature OSNs were further extracted for the second round of subclustering analysis with 1500 anchor features (exclude mitochondrial genes, OR genes, highly-expressed lncRNA Malat1, and sex-specific gene Xist) and the top 20 principal components for clustering at a resolution of 1.

In addition, we downloaded BAM files of previously published raw scRNA-seq data (GSM5283254, GSM5283255, GSM5283256, GSM5283257, GSM5283258, GSM5283259) derived from the main olfactory epithelium of adult mice housed in a typical homecage environment reported by Tsukahara et al (Tsukahara et al., 2021). BAM files were converted back into FASTQ files by bamtofastq executable (https://github.com/10XGenomics/bamtofastq/releases). Next, the published scRNA-seq data were aligned to the *Mus musculus* reference genome mm10 (Ensembl 98) and converted into gene-barcode matrices using 10x Genomics Cell Ranger (Zheng et al., 2017) version 4.0.0. We focused on cells that were regarded as mature OSNs in the original paper. After rigorous quality control, we removed cells from the homecage-4 sample. Next, scRNA-seq data of the remaining mature OSNs from Tsukahara et al (Tsukahara et al., 2021) were integrated with scRNA-seq data of mature OSNs from this study, with 1500 anchor features (excluding mitochondrial genes, OR genes, highly-expressed lncRNA Malat1, and sex-specific gene Xist) and the top 25 principal components for clustering at a resolution of 0.2. A small cluster expressing Ifi27 was identified but not used for further analyses in this paper.

By subclustering analysis, mature OSNs could be divided into two groups: Cd36+ OSNs and Cd36-OSNs. Differentially expressed genes between Cd36+ OSNs and Cd36-OSNs were identified with the Wilcoxon rank-sum test by the FindAllMarkers() function. Only those with an absolute fold change ≥ 1.5 and adjusted p-value < 0.05 were considered as differentially expressed genes. Functional enrichment of differentially expressed genes was performed by clusterProfiler (Yu et al., 2012) version 4.2.2. When calculating the percentage of dorsal and ventral OSNs in each group of OSNs, the OSNs were classified as dorsal or ventral based on their expression of known dorsal marker genes (*Nqo1*, *Acsm*4) and ventral marker genes (*Ncam*2, *Nfix*, *Nfib*).

### Identification of Cd36+ OSN-enriched OR genes

To increase statistical power, we used integrated scRNA-seq data containing mature OSNs from this study and Tsukahara et al (Tsukahara et al., 2021) to identify Cd36+ OSN-enriched OR genes. We first identified the OR gene expressed in each mature OSN from integrated scRNA-seq data. Mature OSNs from Tsukahara et al (Tsukahara et al., 2021) which were found to contradict the previously published annotations of expressed OR genes were removed from this analysis. All cells expressing the same OR gene were considered as an OSN subgroup. 945 OSN subgroups with at least 10 cells in each subgroup were identified and used in this analysis. For each OSN subgroup, we calculated the number of cells that were found within Cd36+ OSNs and Cd36-OSNs and performed a binomial test to determine if the expressed OR gene was biased towards any of the two groups. In the scRNA-seq data analyzed, there were 2573 Cd36+ OSNs and 44829 Cd36- OSNs. If the expressed OR gene was not biased towards any of the two groups, the proportion of cells in this OSN subgroup that were found within Cd36+ OSNs and Cd36-OSNs should be the same as the proportion of cells in Cd36+ OSNs and Cd36-OSNs (2573/44829≈0.057 in our case). We used the Benjamini-Hochberg method to correct multiple testing. OSN subgroups with Benjamini-Hochberg corrected p-value < 0.05 are regarded as biased OSN subgroups. Among them, we distinguished those predominantly composed of Cd36+ OSNs and considered their expressed OR genes as Cd36+ OSNs-enriched OR genes. In contrast, the expressed OR genes of subgroups primarily consisting of Cd36- OSNs were categorized as Cd36+ OSNs-depleted OR genes. To examine if Cd36+ OSNs-enriched OR genes were enriched in a particular chromosome, a binomial test for each chromosome was performed. If Cd36+ OSNs-enriched OR genes were not enriched in the chromosome, the proportion of Cd36+ OSNs-enriched OR genes in this chromosome and all chromosomes should be the same as the proportion of OR genes in this chromosome and all chromosomes.

### Phylogenetic analysis of OR genes

Phylogenetic analysis was performed using 1140 annotated mouse OR genes. We retrieved amino acid sequences of mouse OR genes from the Ensembl database and made a multiple sequence alignment using MUSCLE v5 (Edgar, 2004). Next, we constructed a maximum likelihood phylogenetic tree using RAxML (Stamatakis, 2014) with the automatic selection of amino acid selection model (PROTGAMMAAUTO).

### scATAC-seq data processing and cell type identification

For each scATAC-seq dataset, raw reads were preprocessed using 10x Genomics Cell Ranger ATAC (Satpathy et al., 2019) version 2.0.0, including read filtering, alignment to Mus musculus reference genome mm10 provided by 10X Genomics, peak calling, cell calling, and generating peak-barcode count matrix. The count data and fragment files were loaded into R (Version 4.1.3) and further processed using the R package Siganc (Stuart et al., 2021). For each scATAC-seq data, we calculated the nucleosome signal (the ratio of mononucleosomal to nucleosome-free fragments), TSS enrichment score, and the percentage of reads in peaks for each cell. High-quality cells with 5000-40000 fragments, TSS enrichment score >= 4, nucleosome signal <2, and the percentage of reads in peaks >= 40 were retained for downstream analysis. Next, scATAC-seq datasets across two different time points were integrated using the integration workflow (https://stuartlab.org/signac/articles/integrate_atac.html). First, the peaks called in each dataset were merged using the reduce() function of the GenomicRanges (Lawrence et al., 2013) package to create a common peak set. Next, the count matrix for each dataset was constructed using this common peak set. To reduce dimensions, latent semantic indexing (LSI) was performed for each dataset. LSI involves two steps. The first step is to normalize data using the frequency-inverse document frequency (TF-IDF) transformation by RunTFIDF() function. Subsequently, the features present in more than *20* cells were used to run singular value decomposition (SVD) on the TF-IDF matrix. To integrate the two datasets and remove the batch effect, we identified integration anchors between the two datasets using reciprocal LSI projection by the FindIntegrationAnchors() functions and used these anchors to integrate LSI embeddings across the two datasets using the IntegrateEmbeddings() function. Finally, LSI components 2 to 30 were used to perform uniform manifold approximation and projection (UMAP) dimensional reduction and clustering at a resolution of 0.8. It yielded 22 clusters. To annotate each cluster, the activity of each gene was quantified by assessing the chromatin accessibility associated with each gene and these gene activity scores were normalized. Besides, the pre-processed integrated scRNA-seq dataset was used for cross-modality integration and label transfer, which also helped to interpret the identity of each cluster. By examining both gene activity scores of previously mentioned marker genes and predicted labels, we first annotated cells that do not belong to the olfactory neuronal lineage: sustentacular cells, ensheathing glia, bowman’s gland, periglomerular cells, brush cells, B cells, erythrocytes, basophils, neutrophils, and monocytes. Due to the incapability to distinguish HBCs and GBCs based on the above preliminary clustering, cell clusters that putatively corresponded to cells in the olfactory neuronal lineage (HBCs, GBCs, INPs, Immature OSNs, and mature OSNs) were extracted for further subclustering analysis. Peaks calling was performed using data from cells in the olfactory neuronal lineage using the CallPeak() function. These peaks were used to generate a count matrix for cells in the olfactory neuronal lineage of each dataset. Next, the same integration workflow of Signac as mentioned above was applied to the olfactory neuronal lineage subclustering analysis except for the used features and clustering resolution. We used features that present in more than 10 cells and a clustering resolution of 1.2 for olfactory neuronal lineage subclustering analysis. Similarly, the gene activity matrix was created and normalized. The pre-processed scRNA-seq dataset of olfactory neuronal lineage was used for cross-modality integration and label transfer. By examining both gene activity scores of previously mentioned marker genes and the predicted labels, cells of the olfactory neuronal lineage were precisely annotated: HBCs, GBCs, INPs, immature OSNs, and mature OSNs. Besides, cells that were identified as not belonging to the olfactory neuronal lineage were annotated using their corresponding marker genes.

### Subclustering analysis of mature OSNs scATAC-seq data

Based on the annotations of the integrated scATAC-seq data, mature OSNs were extracted for further subclustering analysis. The Signac integration workflow was applied to the sub-clustering analysis of mature OSNs as mentioned above except for the features and cluster resolution used. For mature OSNs subclustering analysis, features present in more than 20 cells and a clustering resolution of 0.3 was used. Likewise, mature OSNs from scATAC-seq data could be divided into two groups: Cd36+ OSNs and Cd36- OSNs.

Differentially accessible peaks between Cd36+ OSNs and Cd36- OSNs were identified using a logistic regression model with the total number of counts in each cell included as a latent variable by the FindMarkers() function. Only those with absolute fold change > 0 and Bonferroni adjusted p-value < 0.05 were considered as differentially accessible peaks. Hypergeometric tests were used to test for enrichment of each DNA motif in the set of differentially accessible peaks compared to a background set of peaks with matched GC content by the FindMotifs() function. To further identify differentially active motifs between cell types, motif activities for each cell were measured by running ChromVAR. After integrating with mature OSNs scRNA-seq data, peak-to-gene linkage analysis was performed by computing the correlation between gene expression and accessibility at peaks within 500 kb of the transcription start site(TSS), while correcting technical bias, which was implemented in the LinkPeaks() function. To construct the cell type-specific gene regulatory network (GRN), we first selected candidate transcription factors (TFs) based on the results of motif enrichment analysis for differentially accessible peaks, differential ChromVAR analysis and TF expression levels. Specifically, candidate key TFs were chosen based on the following criteria: their motifs exhibited significant enrichment within cell-type-specifically up-regulated peaks (p-value < 0.05 and fold enrichment ≥ 1.5), their ChromVAR activities were significantly up-regulated (adjusted p-value < 0.05 and absolute differences ≥ 0.4), and they displayed positive gene expression in the given cell type. Next, we searched the presence of the candidate TF motifs within the cell-type-specifically up-regulated peaks. Subsequently, we identified potential target genes of these peaks by peak-to-gene linkage analysis. Only positive region–target gene links are kept. Each GRN is visualized by Cytoscape (version 3.9.1) (Shannon et al., 2003).

### RNA velocity analysis of the olfactory neuronal lineage

To better profile the development trajectory of OSNs, we first extracted cells of the olfactory neuronal lineage from a scRNA-seq dataset that was not subjected to fluorescence-activated cell sorting for subclustering analysis. The Seurat standard analysis workflow (https://satijalab.org/seurat/articles/pbmc3k_tutorial.html) was applied to the subclustering analysis of the olfactory neuronal lineage. Briefly, the count matrix was log-normalized using the NormalizeData() function and the top 2000 variable features were identified using FindVariableFeatures() function. Before dimensional reduction, gene expression data were scaled using the ScaleData() function. Next, principal component analysis was performed and the top 40 principal components were used for clustering at a resolution of 0.6 and running the UMAP dimensional reduction. It yielded 10 clusters. The cell identity of each cluster was annotated by the following marker genes: HBCs (*Cebpd*, *Krt5*); GBCs (*Ascl1*, *Kit*); INPs (*Neurod1*, *Sox11*); Immature OSNs (*Gng8*, *Gap43*); Adcy3- OSNs (*Omp*, lack of *Adcy3*); Cd36+ OSNs (*Omp*, *Cd36*); Cd36- OSNs (*Omp*, lack of *Cd36*).

To infer the development trajectory of OSNs, we performed RNA velocity analysis on this olfactory neuronal lineage scRNA-seq subclustering dataset (HBCs were excluded because they could regenerate multiple mature cell lineages). First, spliced and unspliced count matrices were constructed using the velocyto command line (La Manno et al., 2018). It generated a loom file. Secondly, the Seurat object of the olfactory neuronal lineage scRNA-seq was converted to an h5ad file in R. The h5ad file for gene expression data and loom file for spliced and unspliced matrix was loaded into Python (3.8.0) and merged as an AnnData object. The RNA velocity analysis was then performed by the Python package scVelo (Bergen et al., 2020) using the recommended workflow (https://scvelo.readthedocs.io/VelocityBasics/), which consists of four main steps: preprocessing, computing first- and second-order moments, estimating velocities, and constructing a velocity graph. Unless otherwise specified, default parameters were used. Briefly, data were first log-normalized. Next, first- or second-order moments are computed for each cell in its nearest 30 neighbours. RNA velocities were estimated using a stochastic model of transcriptional dynamics. Finally, the velocity graph was computed based on cosine similarity between velocities and potential cell state transitions. The graph was then used to project the velocities onto the low dimensional UMAP embedding and compute a velocity pseudotime. Besides, a PAGA velocity graph was computed and visualized by a directed graph with edges corresponding to the connectivity between two clusters. Based on the velocity graph, there were two trajectories for Cd36+ OSNs and Cd36- OSNs, and the two trajectories branched from Adcy3- OSNs. To identify precursor cells of Cd36+ OSNs in Adcy3- OSNs, we extracted Tshz1 expression in Adcy3- OSNs and used a 90% quantile as the cutoff. Cells with Tshz1 expression levels greater than 90% quantile were assigned as precursor cells of Cd36+ OSNs and the rest of the cells were assigned as precursor cells of Cd36- OSNs.

### Bulk RNA-seq data analysis for wild-type and Lhx2 KO mOSNs

We downloaded raw bulk RNA-seq data of wild-type and Lhx2 KO mOSNs from Gene Expression Omnibus (GEO) under accession number GSE93570 (2017) (Monahan et al., 2017). For each bulk RNA-seq dataset, fastp (Chen et al., 2018) version 0.21.0 was used to remove adapters and low-quality reads. Trimmed reads were aligned to *Mus musculus* reference genome mm10 using STAR version 2.5.2 (Dobin et al., 2013). Uniquely aligning reads with alignment quality above 30 were retained for downstream analysis. Differential gene expression analysis between wild-type and Lhx2 KO mOSNs was performed in R with the DESeq2 package (Love et al., 2014).

## Supplemental figure and Table Legends

**Figure S1.**
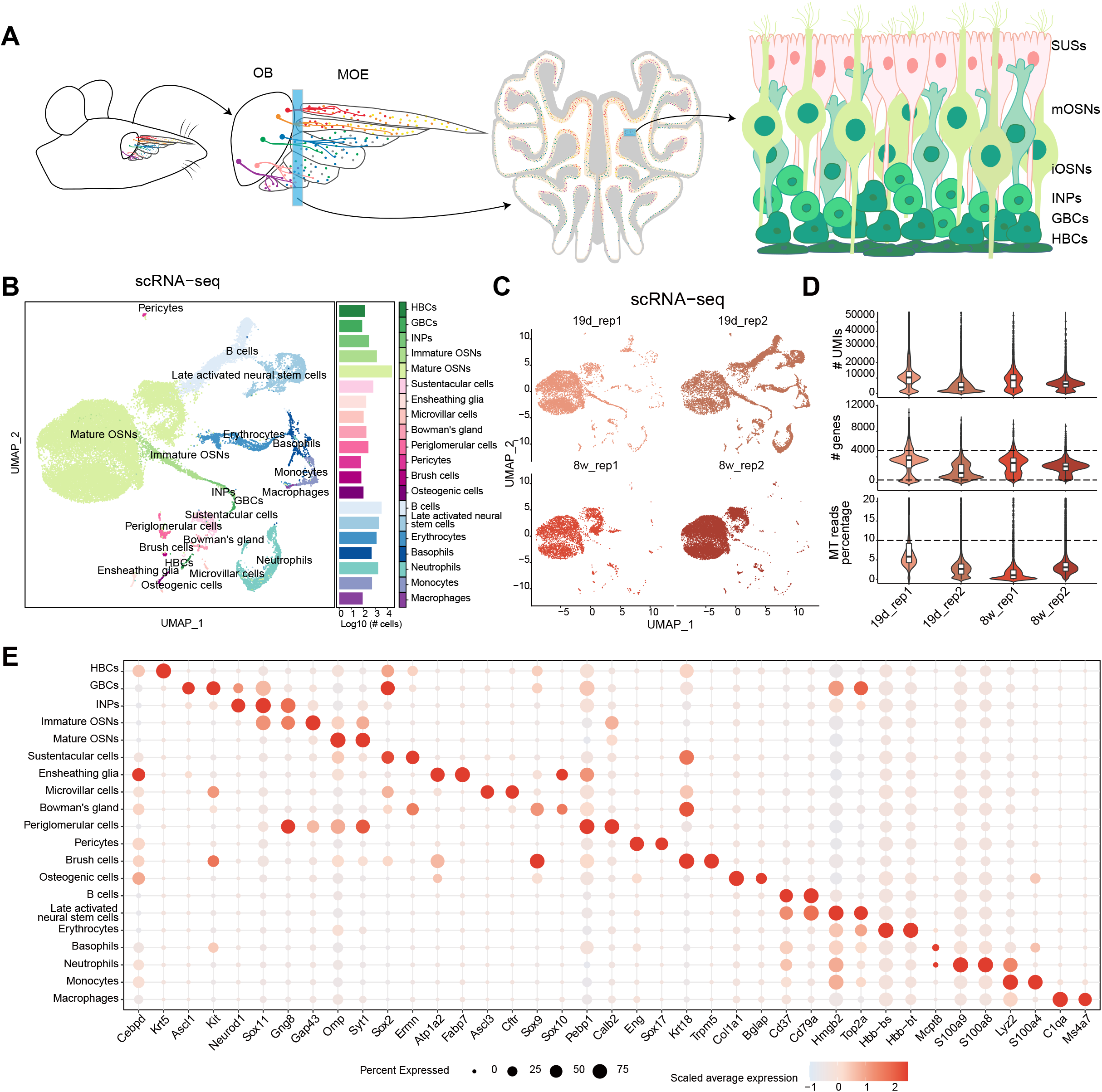
Transcriptome profile of mouse main olfactory epithelium and data quality of scRNA-seq libraries, related to Figure 1. (A) schematic of the anatomy and known major cell types for mouse main olfactory epithelium (MOE). The colorful dots in MOE indicate OSNs and those expressing the same OR gene are shown in the same color. SUCs, sustentacular cells; HBCs, horizontal basal cells; GBCs, globose basal cells; INPs: immediate neuronal precursors; iOSNs: immature olfactory sensory neurons; mOSNs: mature olfactory sensory neurons. (B) Left: uniform manifold approximation and projection (UMAP) plot of the integrated scRNA-seq data from four samples. Each dot represents a single cell colored by its corresponding cell type. Right: bar plot showing the number of cells for each cell type in the scRNA-seq data. Cell-type annotation is labeled to the right of the bar plot. (C) UAMP plots for each scRNA-seq sample. (D) Violin plots showing the distribution of scRNA-seq quality metrics in each sample, including the number of reads, the number of genes, and the percentage of mitochondrial (MT) reads. Cutoffs are indicated by dotted lines. (E) Dot plot showing the expression of marker genes for all cell types annotated in scRNA-seq data. The size of the dot indicates the proportion of cells expressing the gene, and the color of the dot represents the scaled average expression levels.

**Figure S2.**
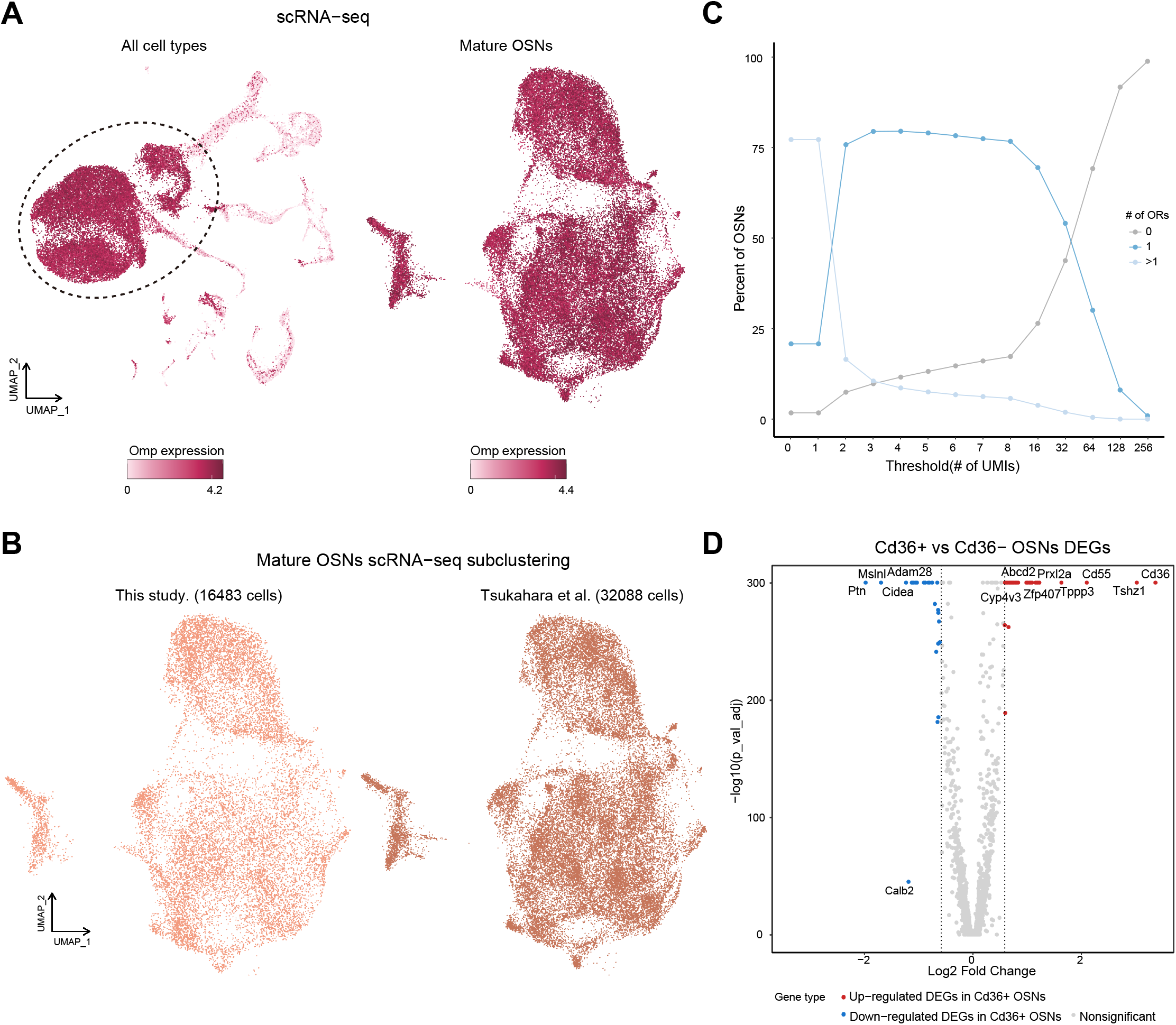
Subclustering analysis of mature OSNs in scRNA-seq data, related to Figure 1. (A) Left: UMAP plot of the integrated scRNA-seq data from four samples, containing all cell types captured by scRNA-seq experiments in this study. Each cell is colored by its expression of Omp, the marker for mature OSNs. Mature OSNs with high expression of Omp are surrounded by a dotted circle and were extracted for further subclustering analysis. Right: UMAP plot of the integrated mature OSNs scRNA-seq data from both this study and Tsukahara et al(Tsukahara et al., 2021). Each cell is colored by its expression of Omp. (B) UAMP plots of the integrated mature OSNs scRNA-seq data from this study (left) and Tsukahara et al (right). The number of cells is annotated above. (C) Percent of OSNs expressing 0, 1, or >1 ORs thresholded by the number of transcripts (**U**nique **M**olecular **I**dentifiers) required for an OR gene to be considered “expressed”. A threshold of 4 UMIs was used and only mature OSNs expressing a single OR gene were retained for downstream analyses. (D) Volcano plot showing differentially expressed genes between Cd36+ OSNs and Cd36- OSNs. The x-axis represents the log2 fold change, and the y-axis represents - log10(p-value_adj). Genes with Bonferroni adjusted-p value < 0.05 and the absolute value of fold change≥1.5 are considered DEGs, while others are nonsignificant DEGs. Red and blue points represent up- and down-regulated DEGs in Cd36+ OSNs, respectively. The grey points represent nonsignificant DEGs.

**Figure S3.**
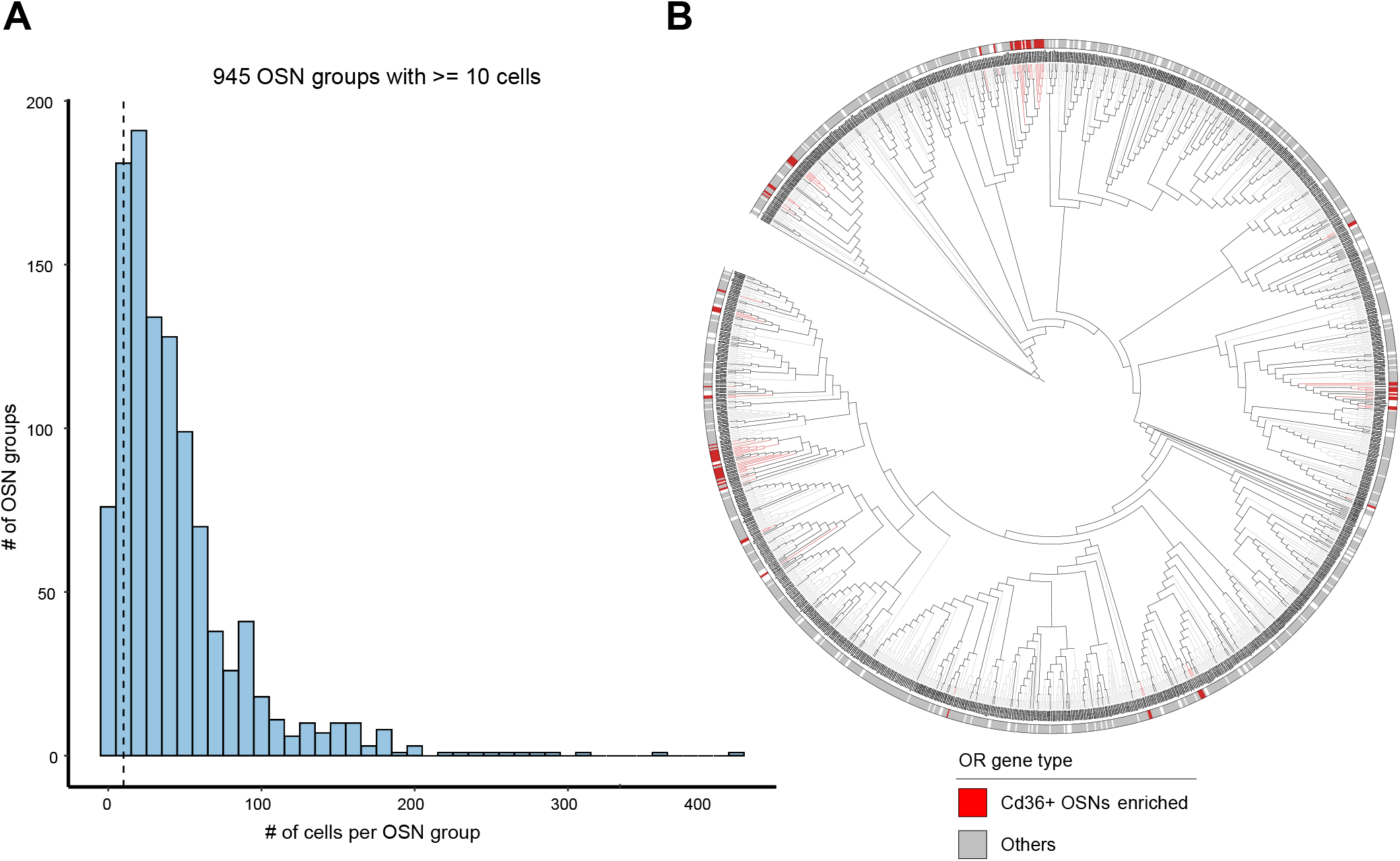
Identification and phylogenetic analysis of Cd36+ OSN enriched ORs, related to Figure 2. (A) Histogram of the number of cells for each OSN group. 945 OSN groups with at least 10 cells in each group were used to identify Cd36+ OSNs-enriched OR genes. (B) Phylogenetic reconstruction of 1140 mouse olfactory receptor proteins. Colored bars at the tree tips indicate if a given OR gene is enriched in Cd36+ OSNs.

**Figure S4.**
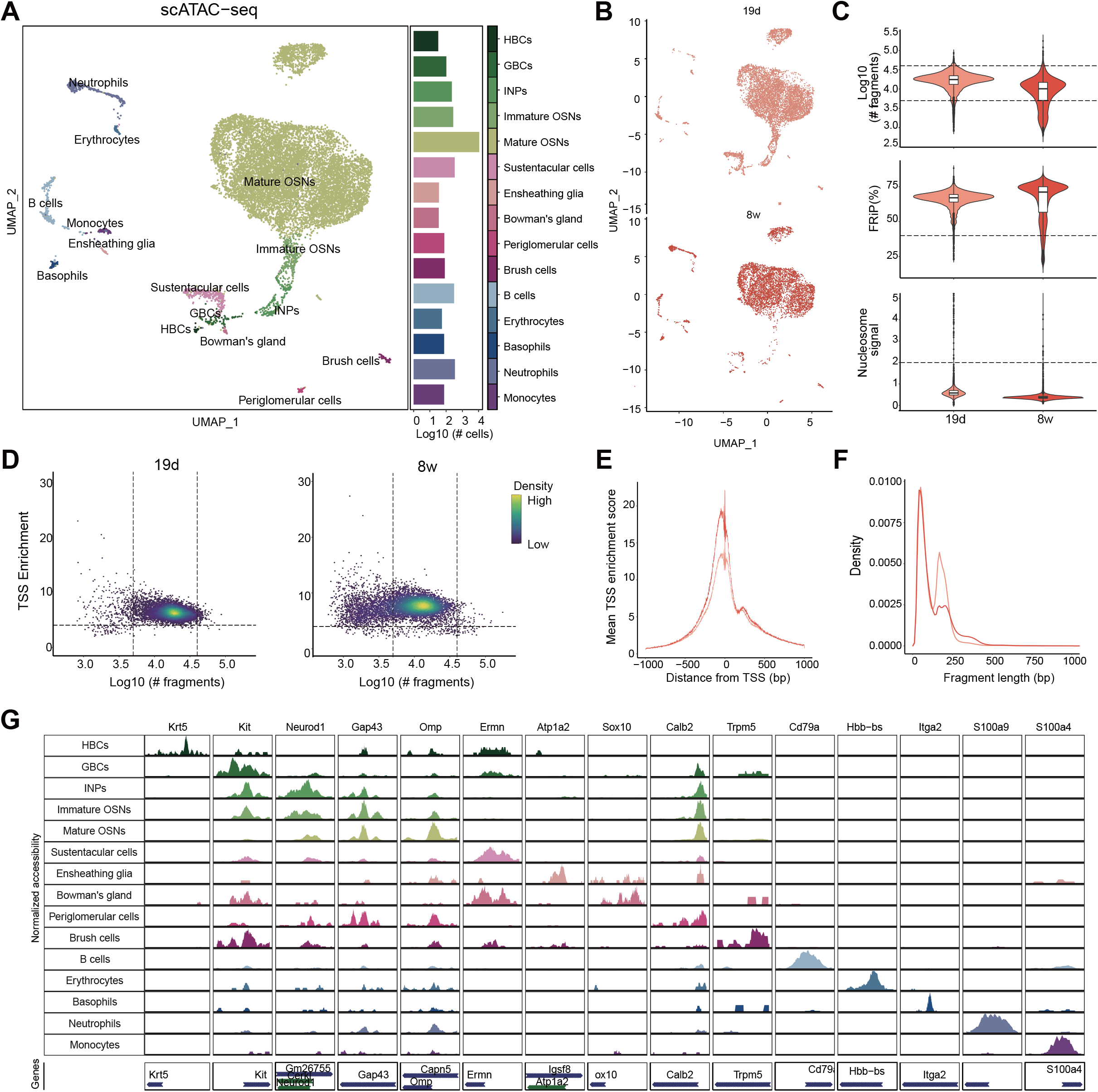
Chromatin profile of mouse main olfactory epithelium and data quality of scATAC-seq libraries, related to Figure 3. (A) Left: UMAP plot of the integrated scATAC-seq data from two samples. Each dot represents a single cell colored by its corresponding cell type. Right: bar plot showing the number of cells for each cell type in the scATAC-seq data. Cell-type annotation is labeled to the right of the bar plot. (B) UMAP plots for each scATAC-seq sample. (C) Violin plots showing the distributions of scATAC-seq quality metrics in each sample, including the number of fragments, the fraction of reads in peaks (FRiP), and nucleosome signal (ratio of mononucleosomal to nucleosome-free fragments). Cutoffs are indicated by dotted lines. (D) scATAC-seq cell thresholding on transcription start site (TSS) enrichment scores and fragment counts for each scATAC-seq sample. The dot color represents the density in arbitrary units of points in the plot. (E) Aggregated normalized accessibility around TSSs for each scATAC-seq sample. (F) Fragment size distributions for each scATAC-seq sample. (G) Genome tracks showing the cell type-specific chromatin accessibility around marker genes for all annotated cell types in scATAC-seq data. The tracks have been normalized to the total number of fragments in each cell type.

**Figure S5.**
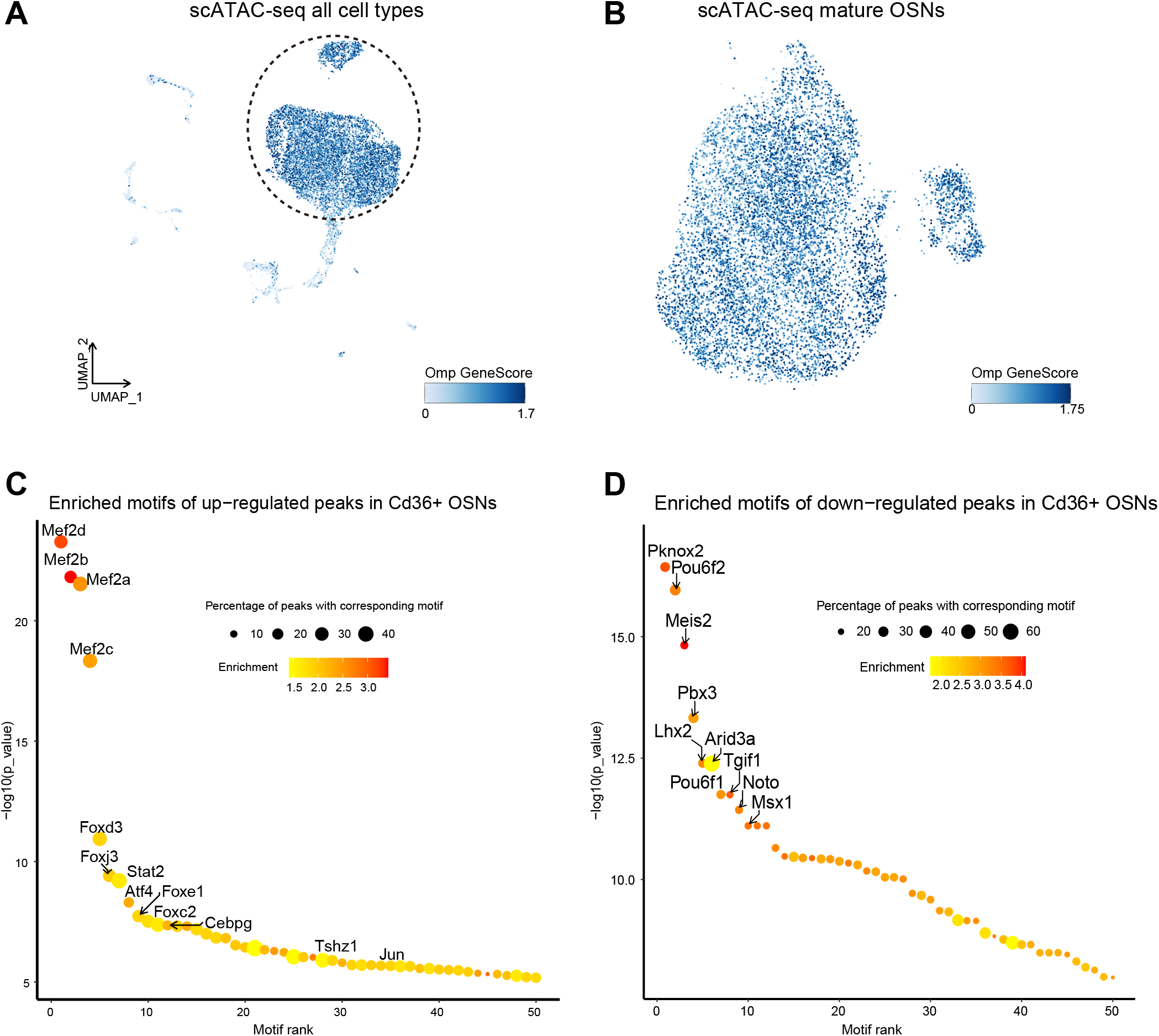
Subclustering analysis of mature OSNs in scATAC-seq data, related to Figure 3. (A) UMAP plot of the integrated scATAC-seq data from two samples, containing all cell types captured by scATAC-seq experiments in this study. Each cell is colored by the gene activity score of Omp, the marker gene for mature OSNs. The dotted circle shows the mature OSNs with high Omp accessibility which were extracted for further subclustering analysis. (B) UMAP plot of the integrated mature OSNs scATAC-seq data. Each cell is colored by the gene activity score of Omp. (C) Dot plot showing enriched motifs in up-regulated peaks of Cd36+ OSNs. The x-axis represents the rank of enriched motif ordered by p-value and the y-axis represents -log10 (p-value). Dot size represents the percentage of target peaks with corresponding motif and dot color represents motif enrichment fold in target peaks compared to background sequences. (D) Dot plot showing enriched motifs in down-regulated peaks of Cd36+ OSNs.

**Figure S6.**
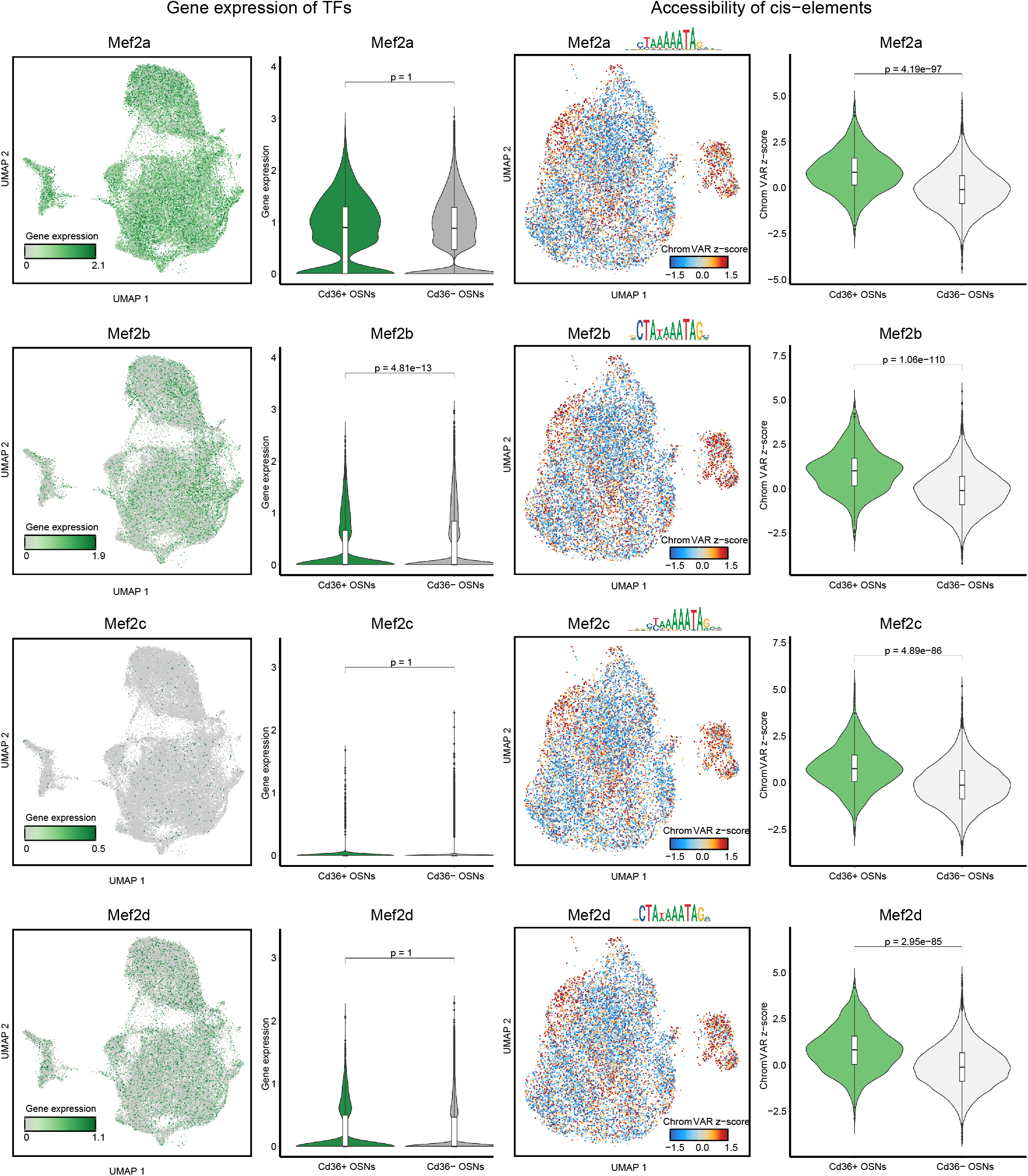
Gene expression and chromVAR z-scores for Cd36+ OSNs enriched TFs, related to Figure 3. (A-D) Left: UMAP plots and violin plots showing the gene expression of TFs from scRNA-seq data. In violin plots, gene expression levels between Cd36+ OSNs and Cd36- OSNs are compared by the Wilcoxon Rank Sum test and the Bonferroni adjusted p-value is annotated above. Right: UMAP plots and violin plots showing chromVAR z-scores for TFs from scATAC-seq data. In violin plots, the chromVAR z-scores of the motif between Cd36+ OSNs and Cd36- OSNs are compared and the adjusted p-value is annotated above.

**Figure S7.**
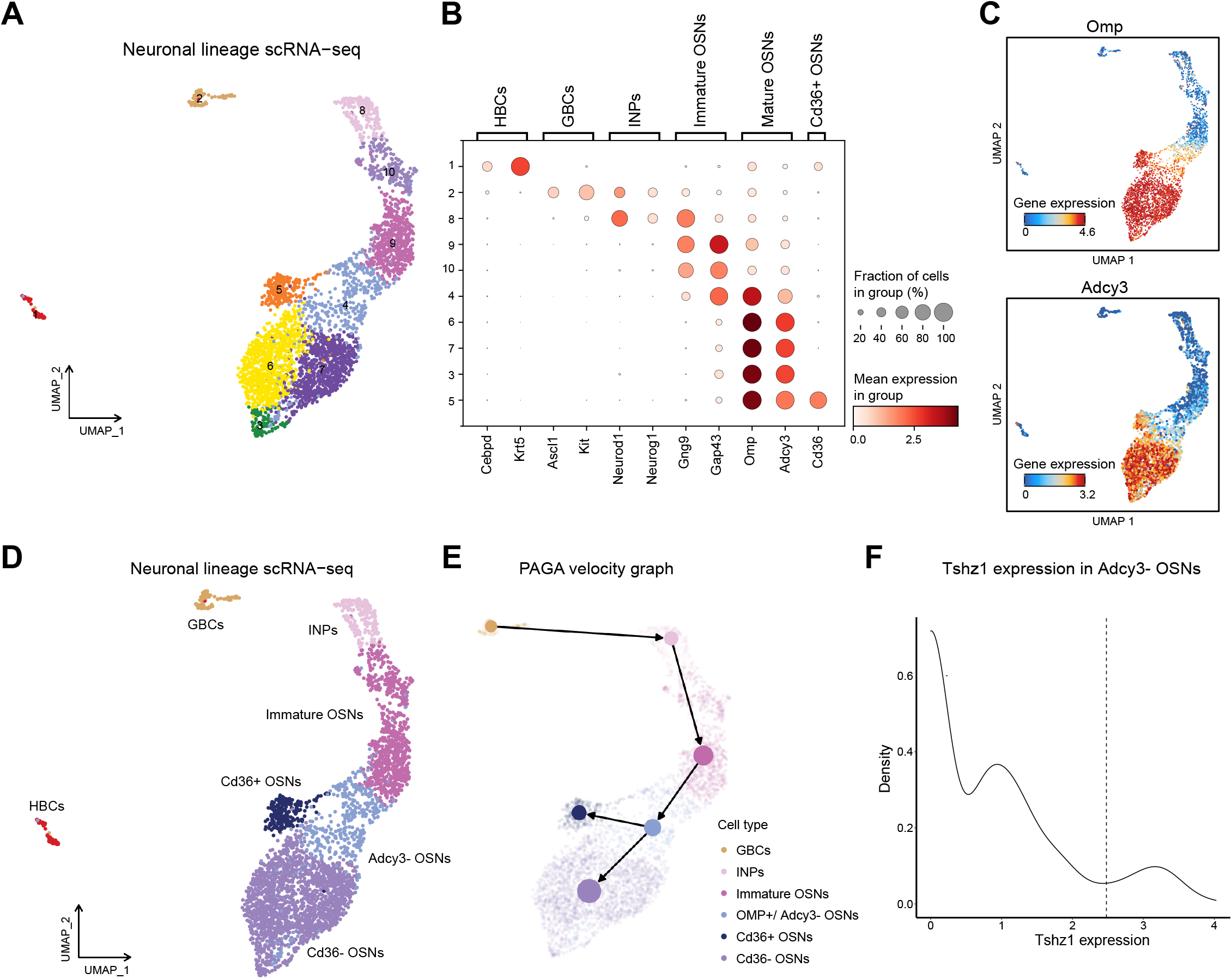
The development of Cd36+ OSNs and Cd36- OSNs, related to Figure 4. (A) UMAP plot showing the subclustering result of olfactory neuronal lineage scRNA-seq data. Each dot represents a single cell colored by its cluster. (B) Dot plot showing expression of cell type specific markers in each cluster identified in Figure S7a. (C) UAMP plots showing expression of Omp (top) and Adcy3 (bottom). (D) UMAP plot showing the subclustering result of olfactory neuronal lineage scRNA-seq data. Each dot represents a single cell colored by its corresponding cell type. (E) An RNA velocity-directed graph, computed by PAGA, projected into the UMAP embedding of olfactory neuronal lineage (HBCs are excluded). Solid lines with arrowheads indicate the directions in which cellular transitions occur. (F) Density plot showing the distribution of Tshz1 expression in Adcy3- OSNs. As shown by the dotted line, cells whose Tshz1 expression exceeds the 90% quantile are considered the precursors of Cd36+ OSNs.

**Figure S8.**
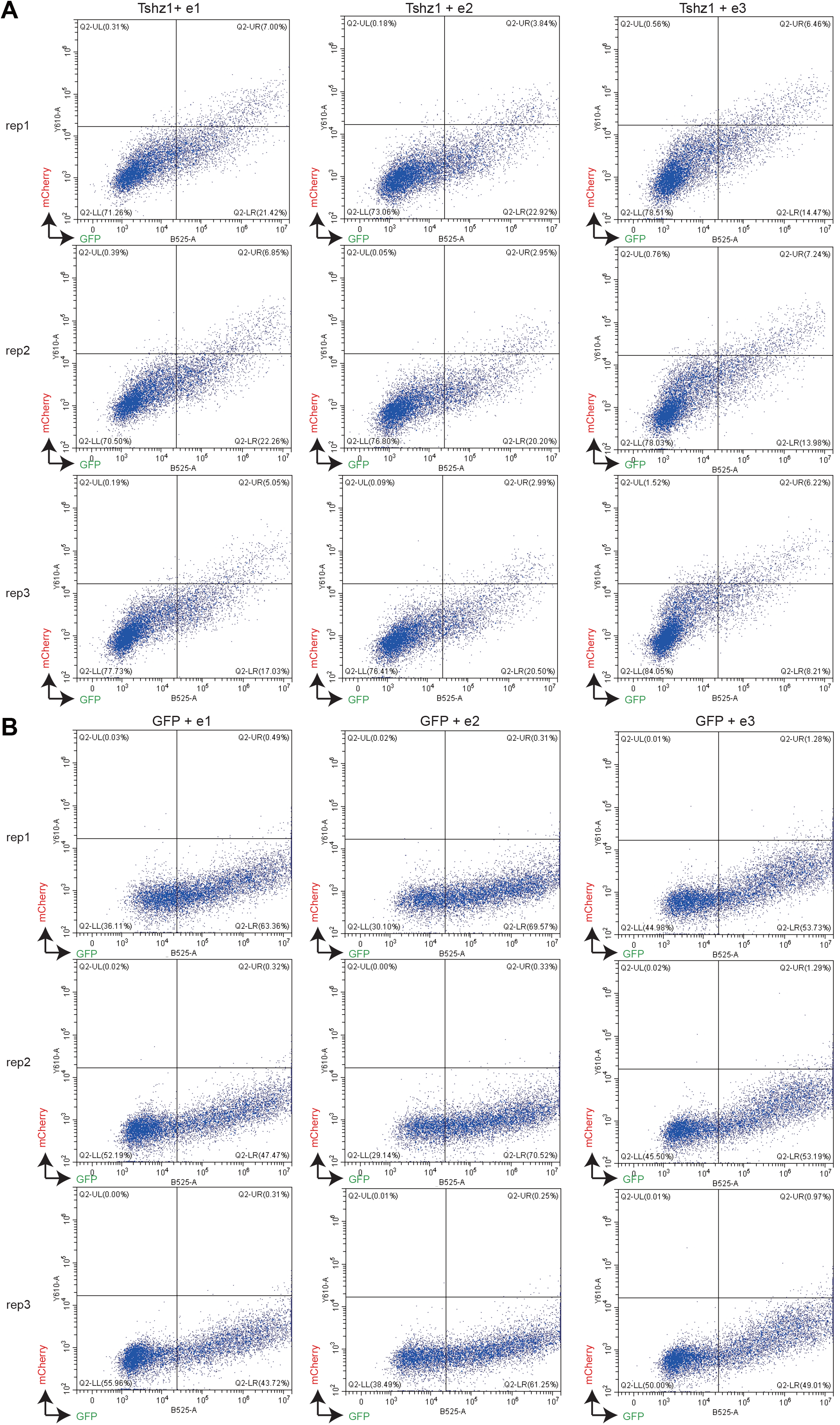
Co-transfection experiments visualized by FACS analysis, related to Figure 6. (A) Two-plasmid co-transfection of N2a cells with Tshz1-GFP and Cd36 enhancer (e1-e3)-mCherry (experimental group). Fluorescence signal visualized by flow cytometer. Each experiment was repeated three times. (B) Two-plasmid co-transfection of N2a cells with GFP and Cd36 enhancer (e1-e3)-mCherry (control group).

**Figure S9.**
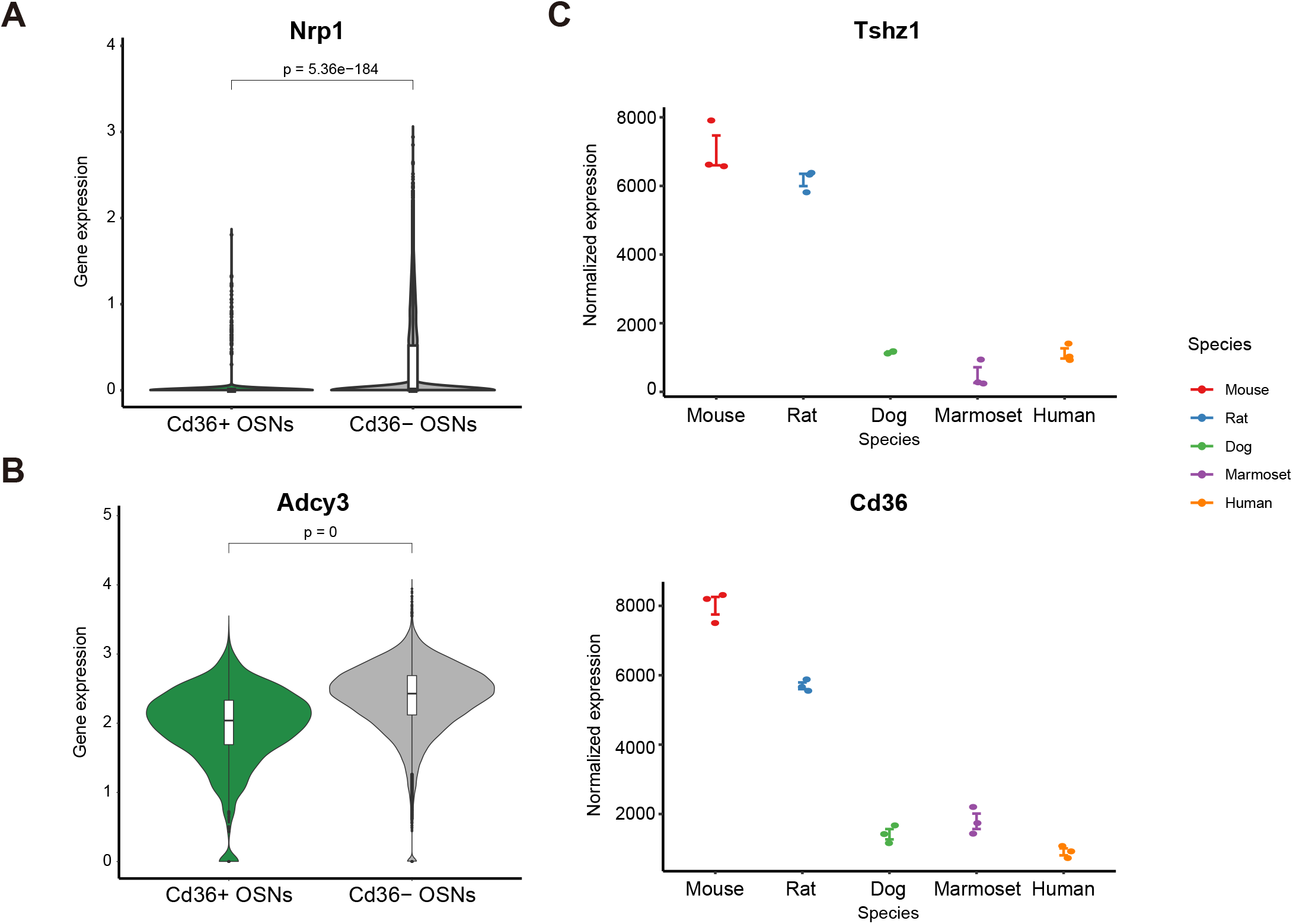
Expression of genes in Cd36+ OSNs and Cd36- OSNs or between species. (A-B) Violin plots showing gene expression levels for (A) Nrp1 and (B) Adcy3 in Cd36+ OSNs and Cd36- OSNs. (C) Normalized gene expression of Tshz1 (top) and Cd36 (bottom) in the olfactory epithelium in five different species obtained from Saraiva LR et al (Saraiva et al., 2019). Each point represents a sample and bars represent standard error.

**Table S1 (related to Figure 1):**
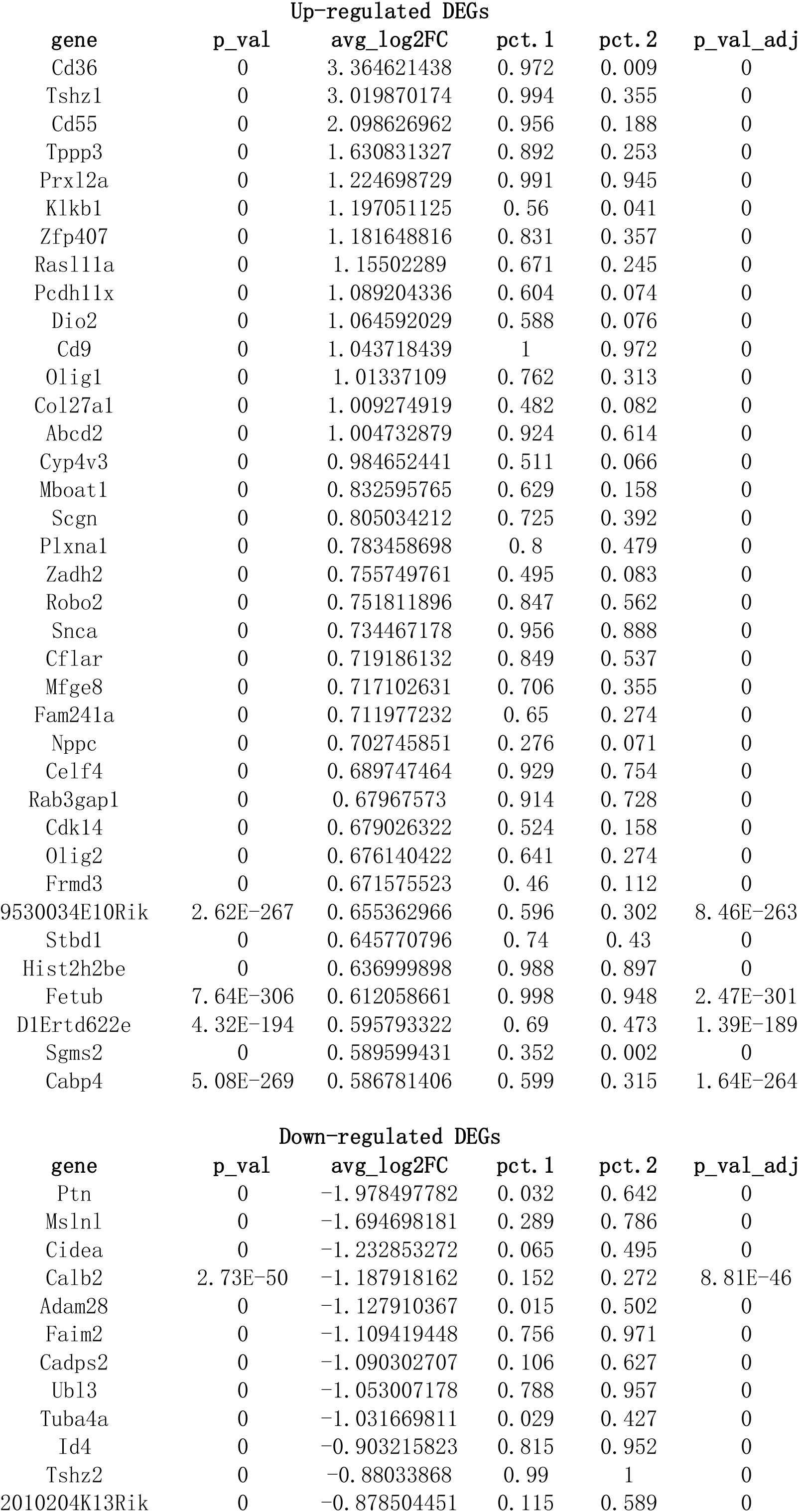
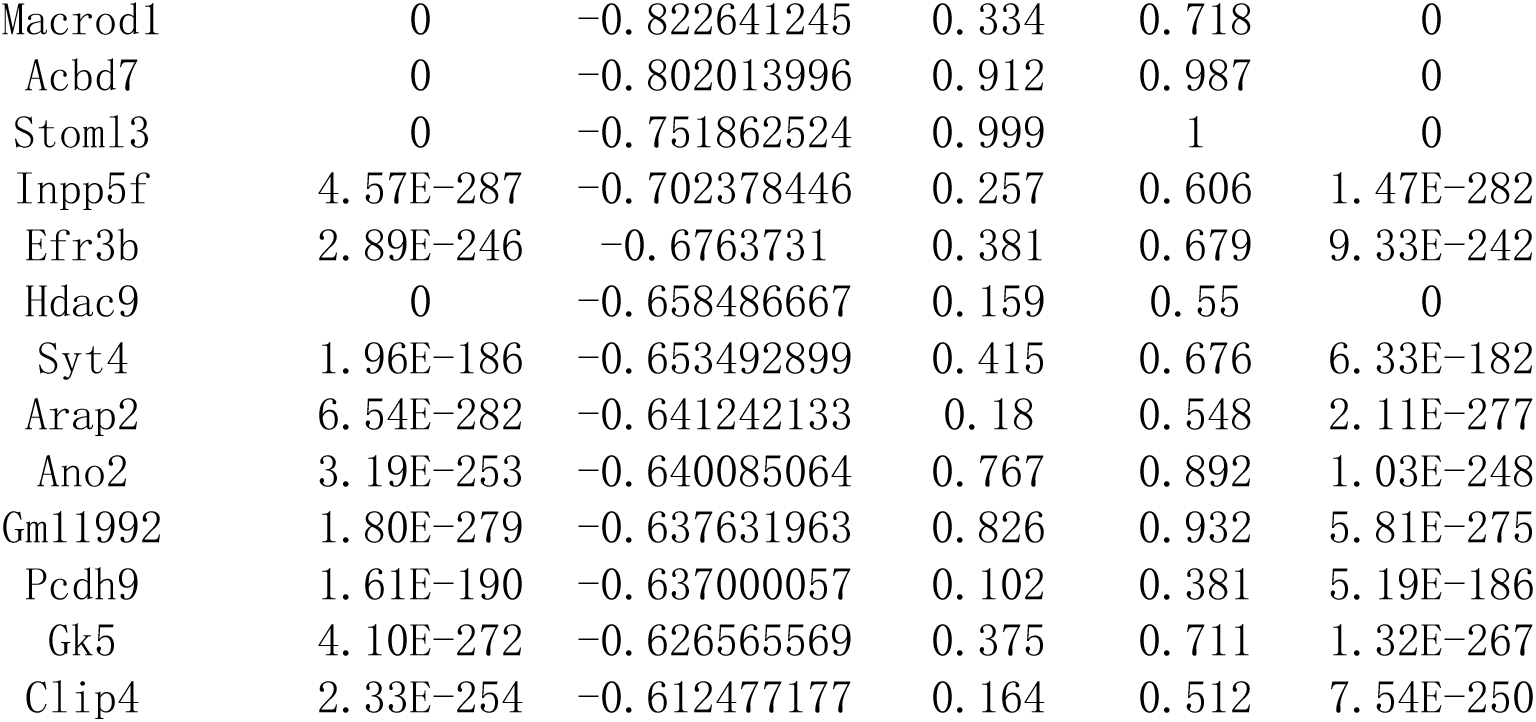
Complete list of up-regulated differentially expressed genes (DEGs) and down-regulated DEGs in Cd36+ OSNs compared to Cd36- OSNs (considering absolute fold change ≥ 1.5 and Bonferroni adjust p-value < 0.05 as cutoff for DEGs).

**Table S2 (related to Figure 3):**
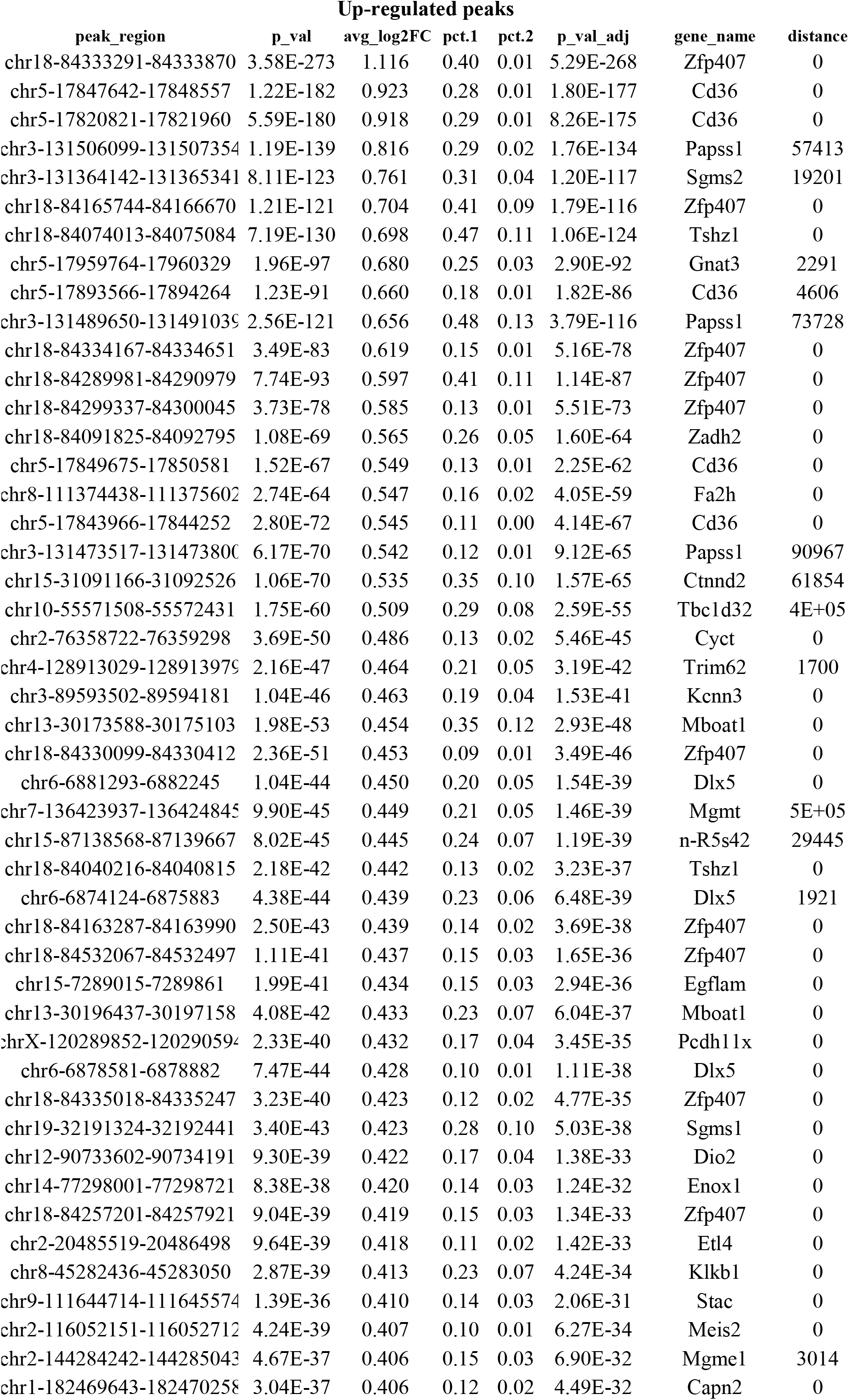
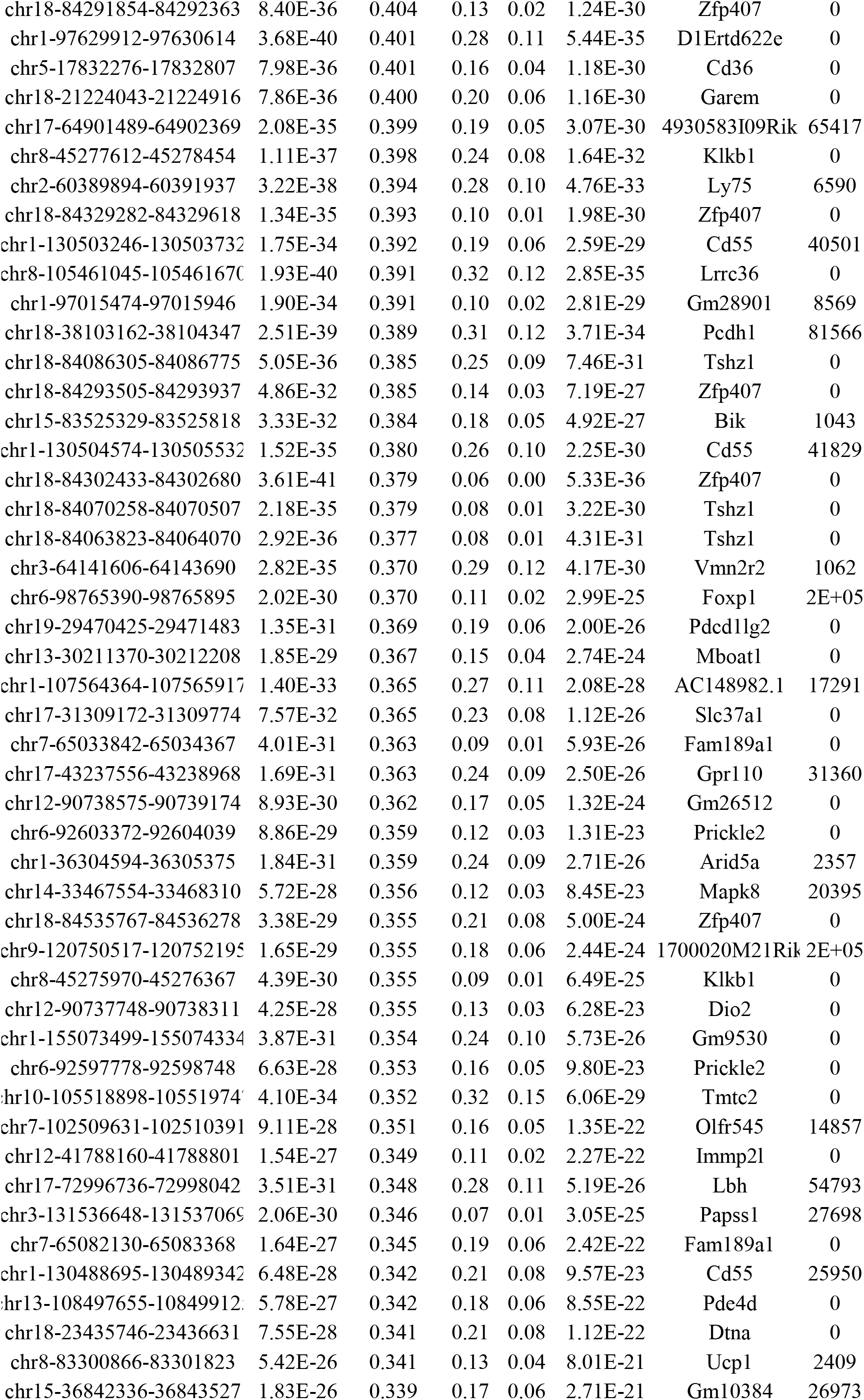
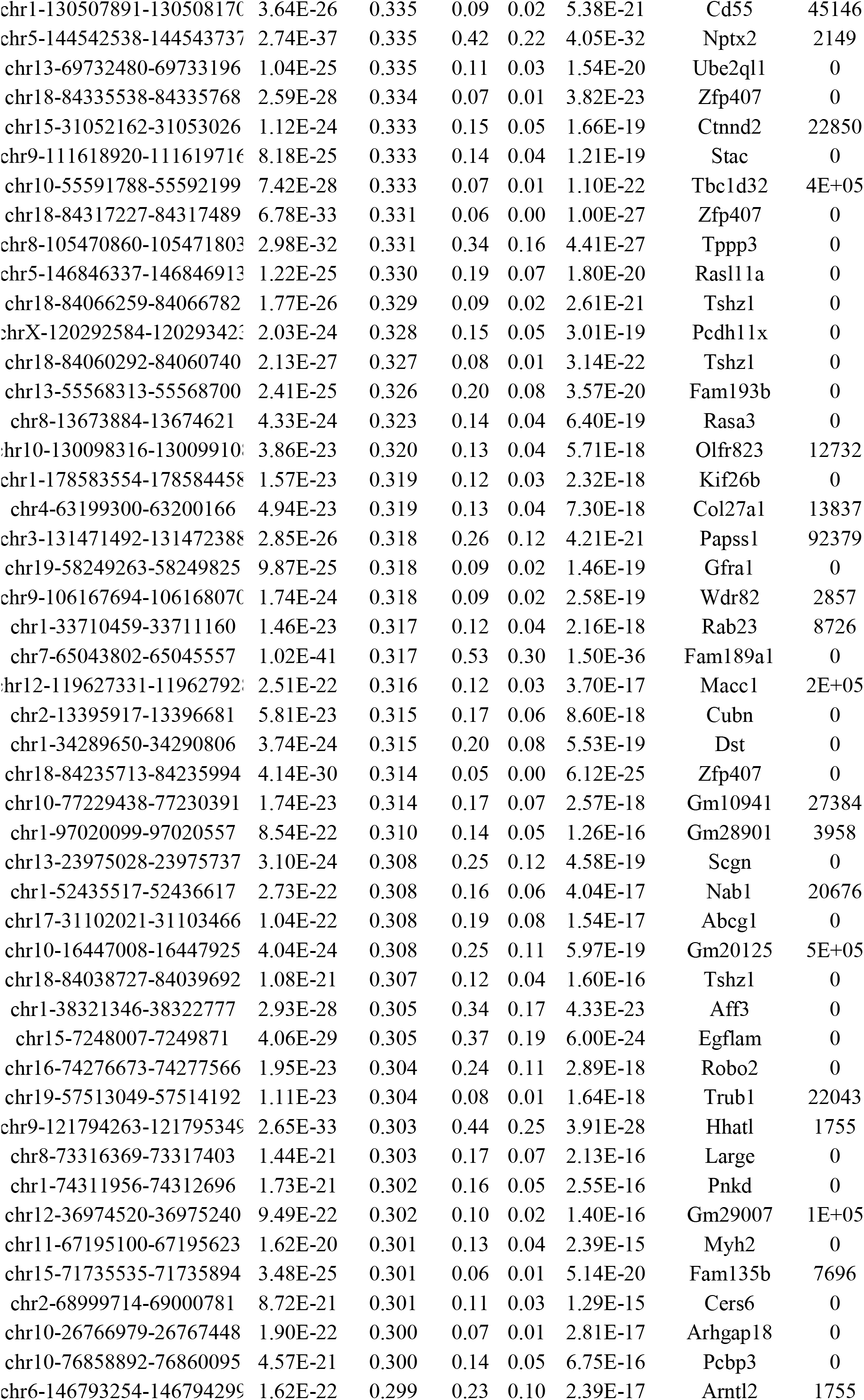
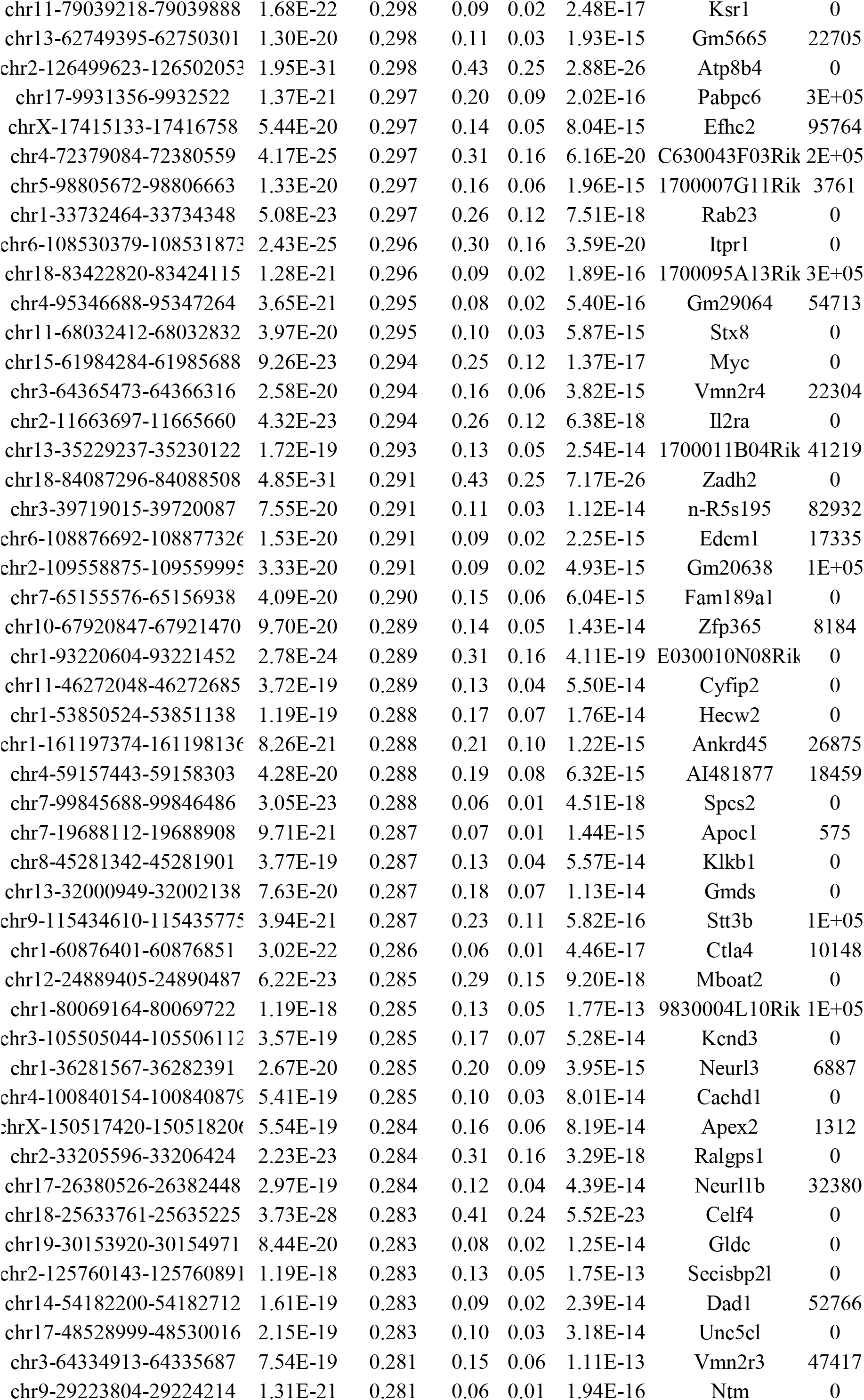
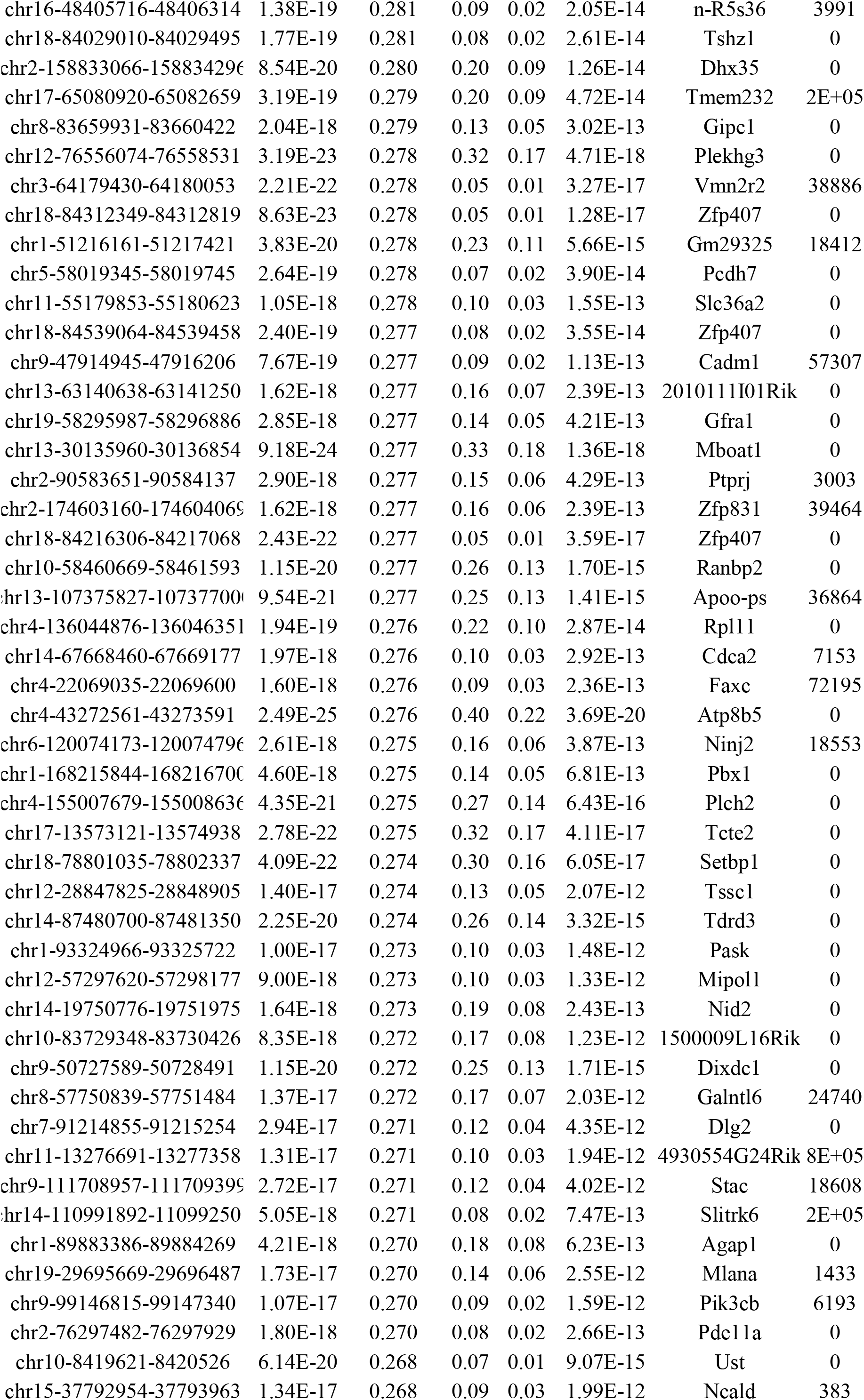
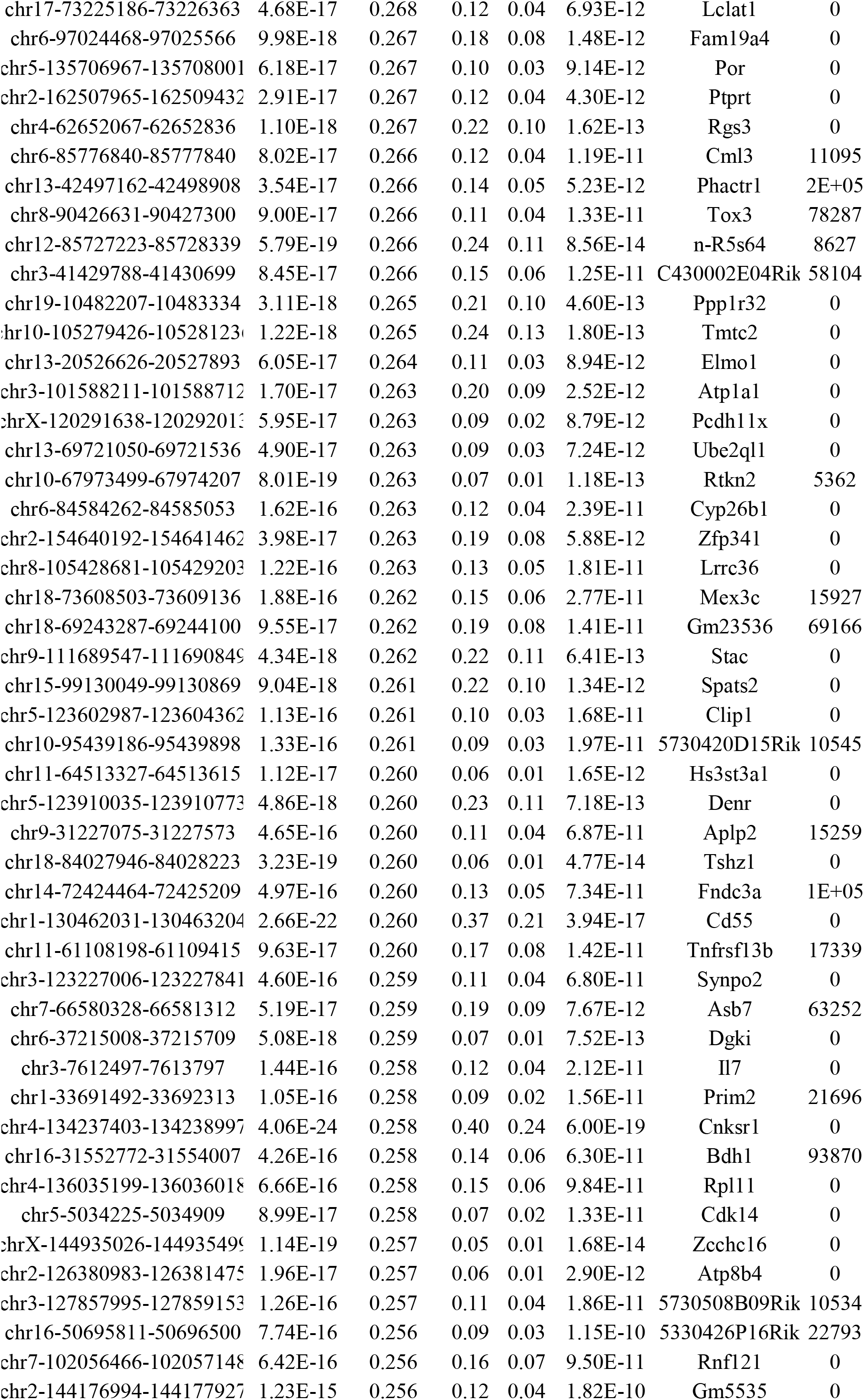
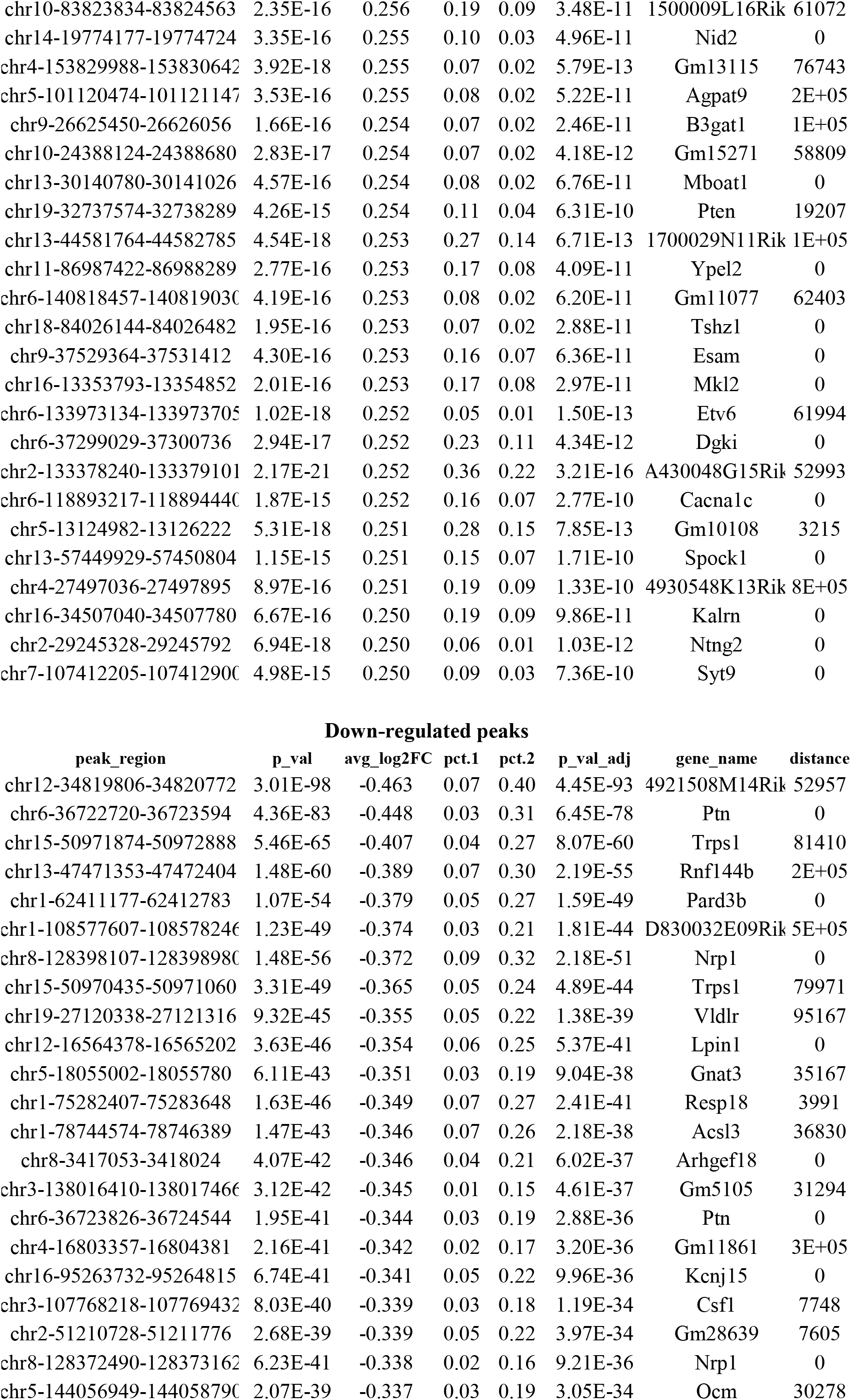
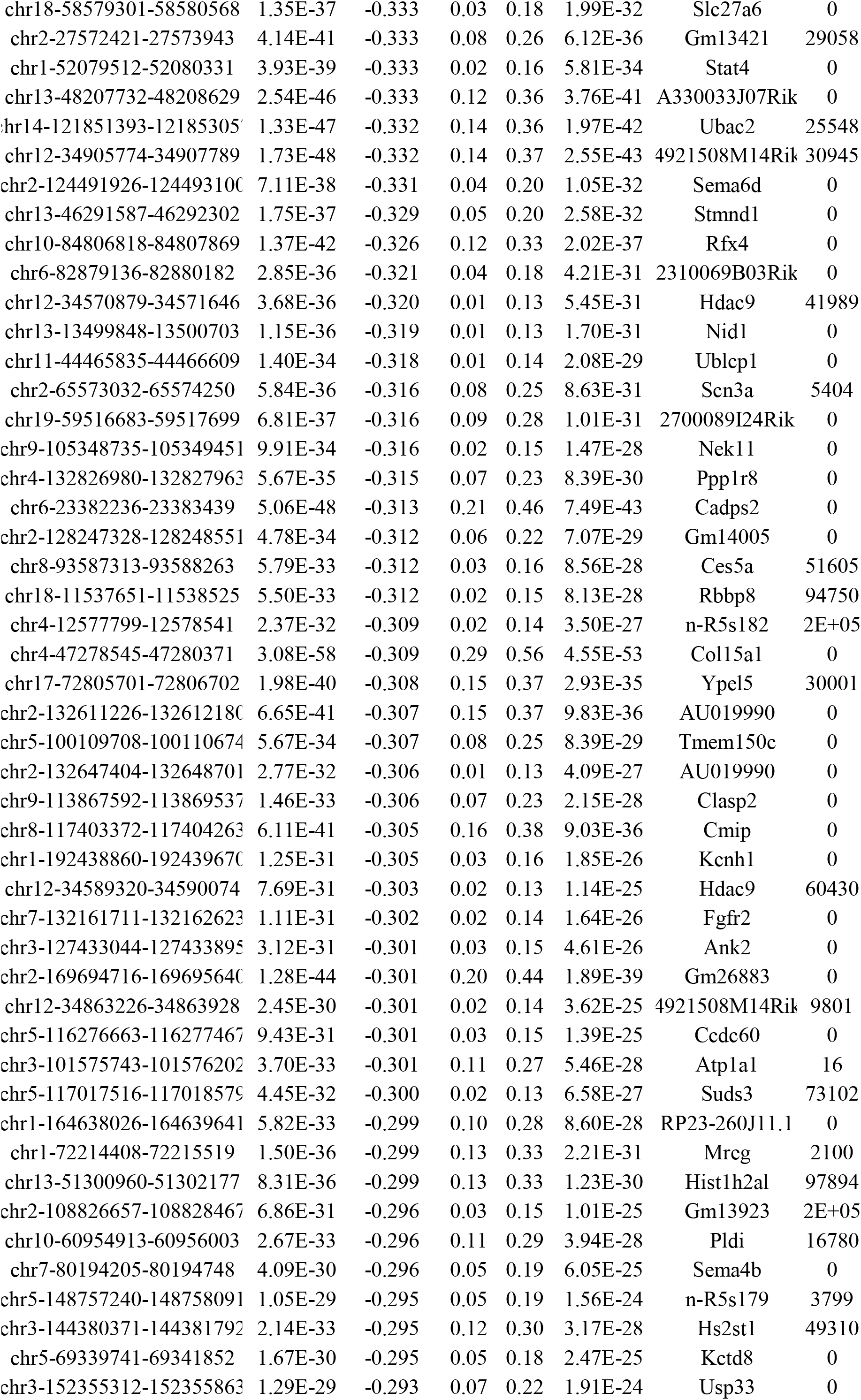
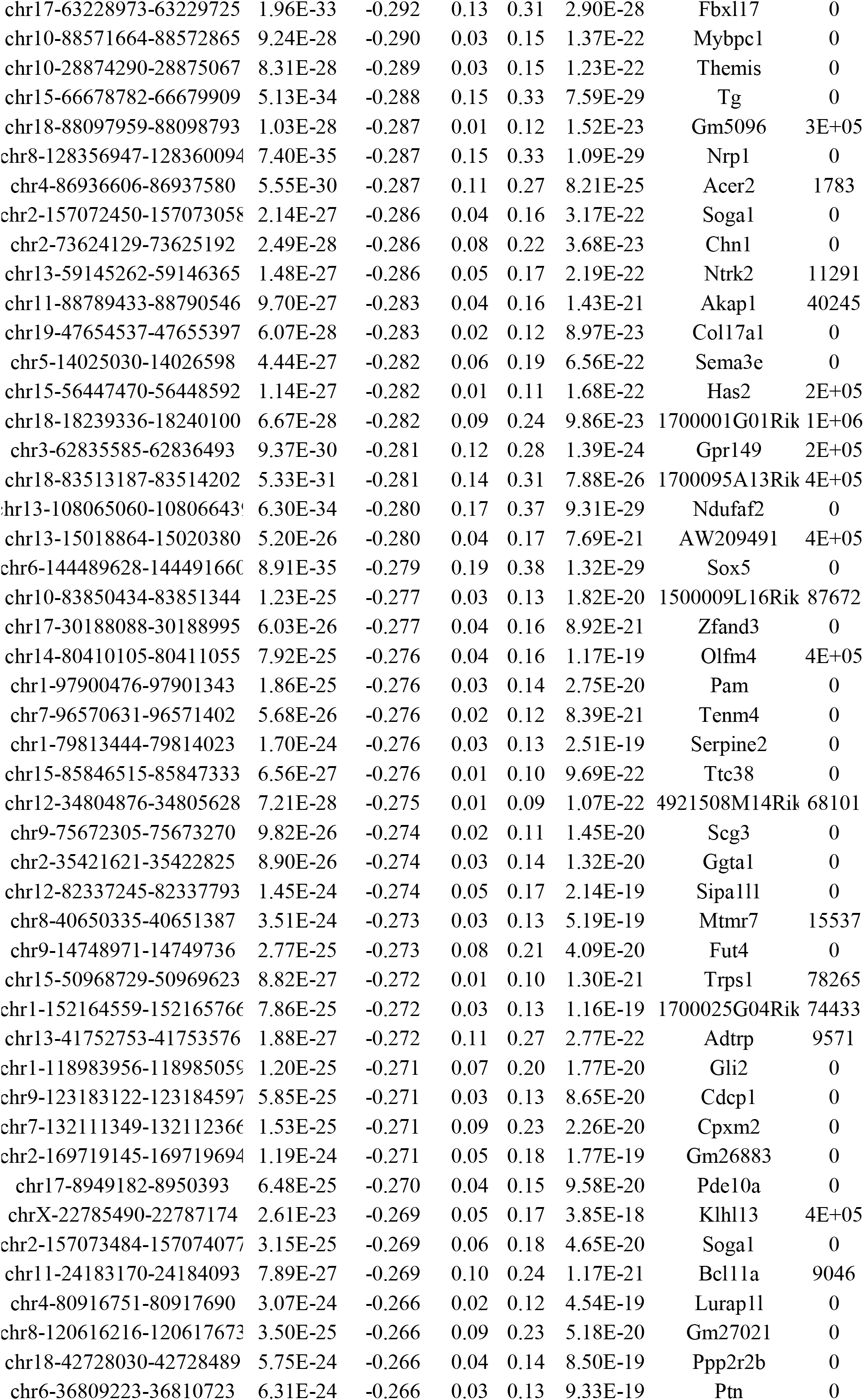
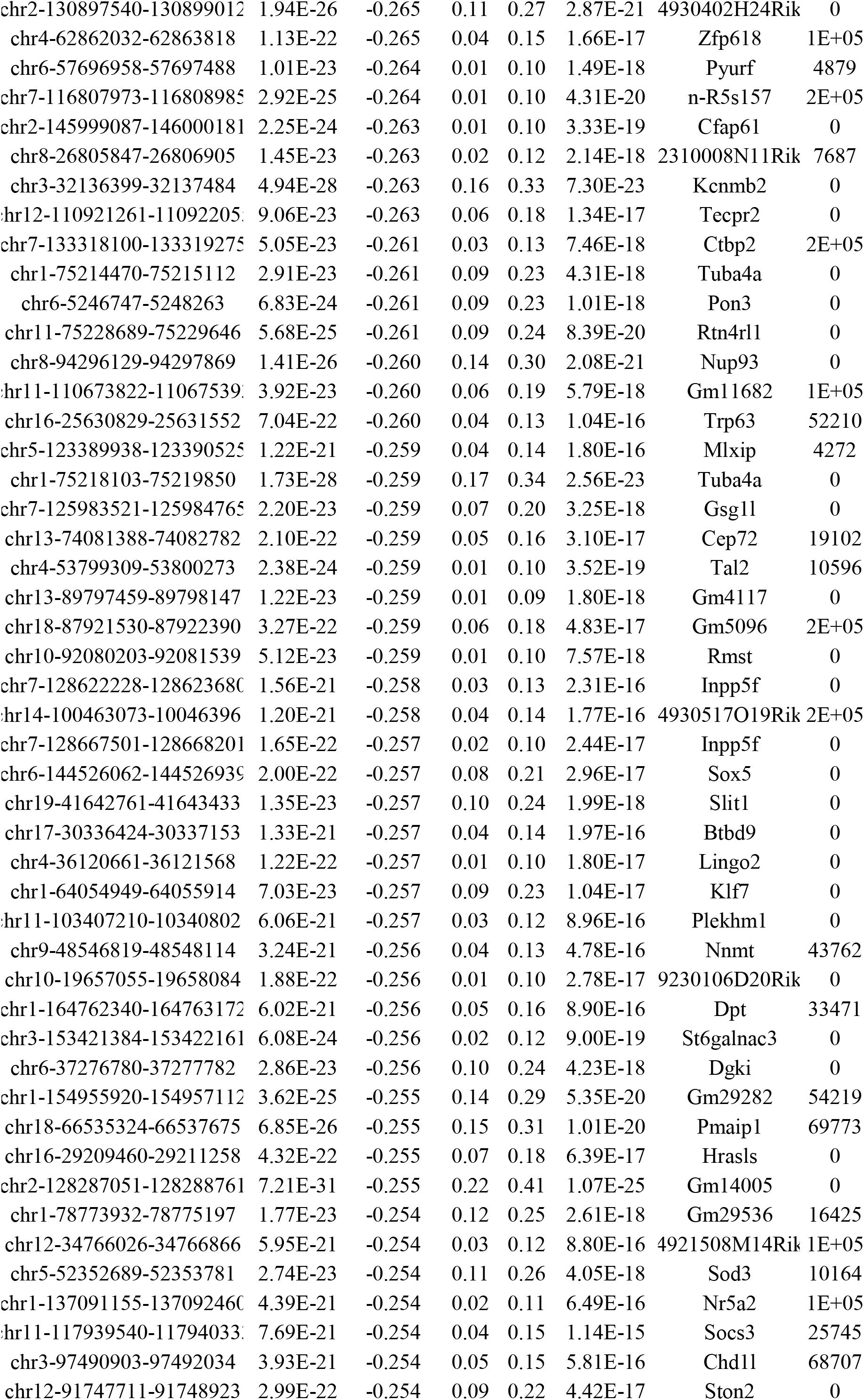
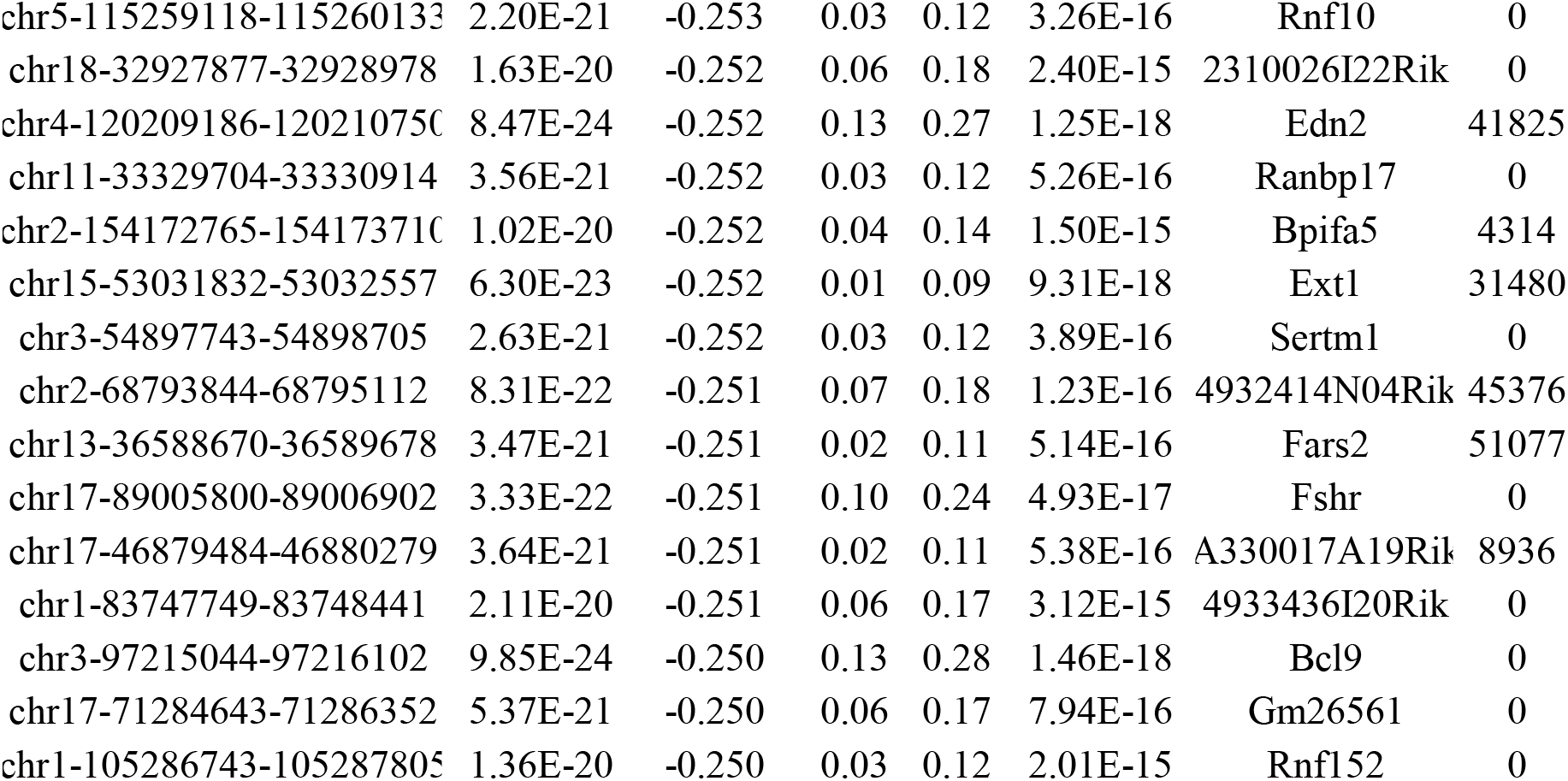
Complete list of up-regulated differentially accessible peaks (DARs) and down-regulated DARs in Cd36+ OSNs compared to Cd36- OSNs (considering absolute fold change > 0 and Bonferroni adjust p-value < 0.05 as cutoff for DARs).

